# A right-hemispheric language network at single-neuron resolution

**DOI:** 10.64898/2026.03.24.713937

**Authors:** Laura F. Schiffl, Lisa M. Held, Felix Waitzmann, Manuel Eder, Hongbiao Chen, Göktuğ Alkan, Paolo Favero, Alexander Utzschmid, Viktor M. Eisenkolb, Moritz Grosse-Wentrup, Julijana Gjorgjieva, Arthur Wagner, Jens Gempt, Bernhard Meyer, Simon N. Jacob

**Affiliations:** Translational Neurotechnology, Department of Neurosurgery, TUM University Hospital, School of Medicine and Health, Technical University of Munich, Germany; Institute for Neuroscience, School of Medicine and Health, Technical University of Munich, Germany; School of Life Sciences, Technical University of Munich, Germany; Munich Cluster for Systems Neurology (SyNergy), Munich, Germany; Research Group Neuroinformatics, Faculty of Computer Science, University of Vienna, Austria; Department of Neurosurgery, TUM University Hospital, School of Medicine and Health, Technical University of Munich, Germany

## Abstract

Human language depends on highly specialized left-hemispheric brain networks. Damage to these networks causes severe language impairments (aphasia), one of the most common, debilitating and costly consequences of left-hemispheric brain injury, especially stroke. The limited recovery of aphasia despite intensive rehabilitation efforts emphasizes the need to understand the basis of residual language abilities at the single-neuron level, which has remained unexplored so far. Here, we report large-scale microelectrode recordings with single-unit resolution over a period of ten months from the right-hemispheric prefrontal and parietal association cortex of an individual with stroke-induced chronic non-fluent aphasia. Single neurons exhibited regionally specific responses during comprehension, retrieval and articulation of words, the core operations of language. Distinct subpopulations encoded linguistic information in a task-specific manner, despite correlated firing patterns across tasks. Both single-neuron activity and temporally coordinated population dynamics were predicted by semantic and phonological embeddings derived from large language models (LLMs), revealing a regional dissociation in which semantic features preferentially accounted for prefrontal activity and phonological features for parietal activity. Our findings suggest that right-hemispheric circuits, homotopic to the left language network, can support language processing through structured, functionally organised activity at the level of single neurons. This study opens an avenue for developing mechanistically specific neurorehabilitation and neurorestorative strategies for aphasia, such as brain-computer interfaces (BCIs), that leverage right-hemispheric language resources.

## Main

Human language relies on a distributed brain network that derives meaning from spoken, written, and gestural signals during comprehension and transforms abstract semantic representations into speech during production^1,2^. Despite extensive work at the systems level, little is known about the neuronal basis of these core language operations. Intracranial recordings in neurological patients have recently demonstrated that individual neurons in the language-dominant fronto-temporo-parietal cortex and hippocampus encode word meaning^3,4^ and participate in speech perception^5–7^ and production^8–11^, providing the first direct evidence that human neurons track semantic information, phonological structure and articulatory plans. However, clinical constraints severely limit the temporal duration and spatial coverage of such recordings, preventing a complete understanding of the single-neuronal and neuronal-network-level organisation of language. This limitation is particularly relevant in the context of stroke-induced aphasia, where linguistic abilities can partially persist after brain injury and recover despite permanent damage to left-hemispheric networks^12–16^. Functional imaging studies have reported language-related activation in both perilesional left-hemispheric cortex and in homotopic regions of the right hemisphere after stroke^17,18^. Such right-hemispheric responses are often discussed as beneficial and compensatory, particularly after extensive left-hemispheric damage^19–23^. Another line of research, however, links their presence to poorer language outcomes and incomplete recovery^24–26^. These opposing views raise a fundamental question: when the left language network is compromised, does the right hemisphere encode structured, task- and content-specific linguistic representations, or does it exhibit non-specific activation patterns that are unlikely to support language processing?

A deeper understanding of the linguistic selectivity of individual neurons in aphasia has profound implications beyond fundamental neuroscience. Recent advances in BCI technology have demonstrated that decoding motor cortical signals can restore communication in patients with speech impairments^27–36^. However, these approaches rely on preserved vocal motor planning. They cannot be translated to aphasia, in which the breakdown occurs earlier in the language processing stream and affects higher-order linguistic representations, such as semantic and phonological information, which are broadly distributed across the cortex^1,37^ and are still poorly characterized at the neuronal level.

Here, we report long-term intracortical microelectrode recordings in an individual with aphasia from four right-hemispheric regions that, on the left, support key language functions in the semantic, phonological, and syntactic domains^2,38–40^: the angular gyrus (ANG), supramarginal gyrus (SMG), inferior frontal gyrus (IFG), and middle frontal gyrus (MFG). Over ten months, the participant performed word repetition, comprehension, and naming tasks, enabling us to examine the extent to which individual neurons contralateral to the canonical left language network support language under different processing demands and over extended time periods.

### Language task performance

We studied an individual with chronic non-fluent (Broca’s) aphasia following an extensive left-hemispheric stroke six years prior to enrollment (**Fig. 1a**). Across 54 sessions spanning ten months, the participant performed three word-level tasks targeting core linguistic processes: word retrieval or naming (NAM), in which an image was visually presented and the participant vocalized the corresponding word; word repetition (REP), in which a spoken word was auditorily presented and the participant repeated it; and word comprehension (COM), in which a spoken word was auditorily presented and the participant identified the matching image in a response display by eye fixation (**Fig. 1b, c**). Henceforth, we refer to the visually or auditorily presented first stimulus in the trial as the sample, and to the visual image matching the auditory sample in the comprehension task, appearing as one of three sequentially presented stimuli, as the target.

**Fig. 1.**
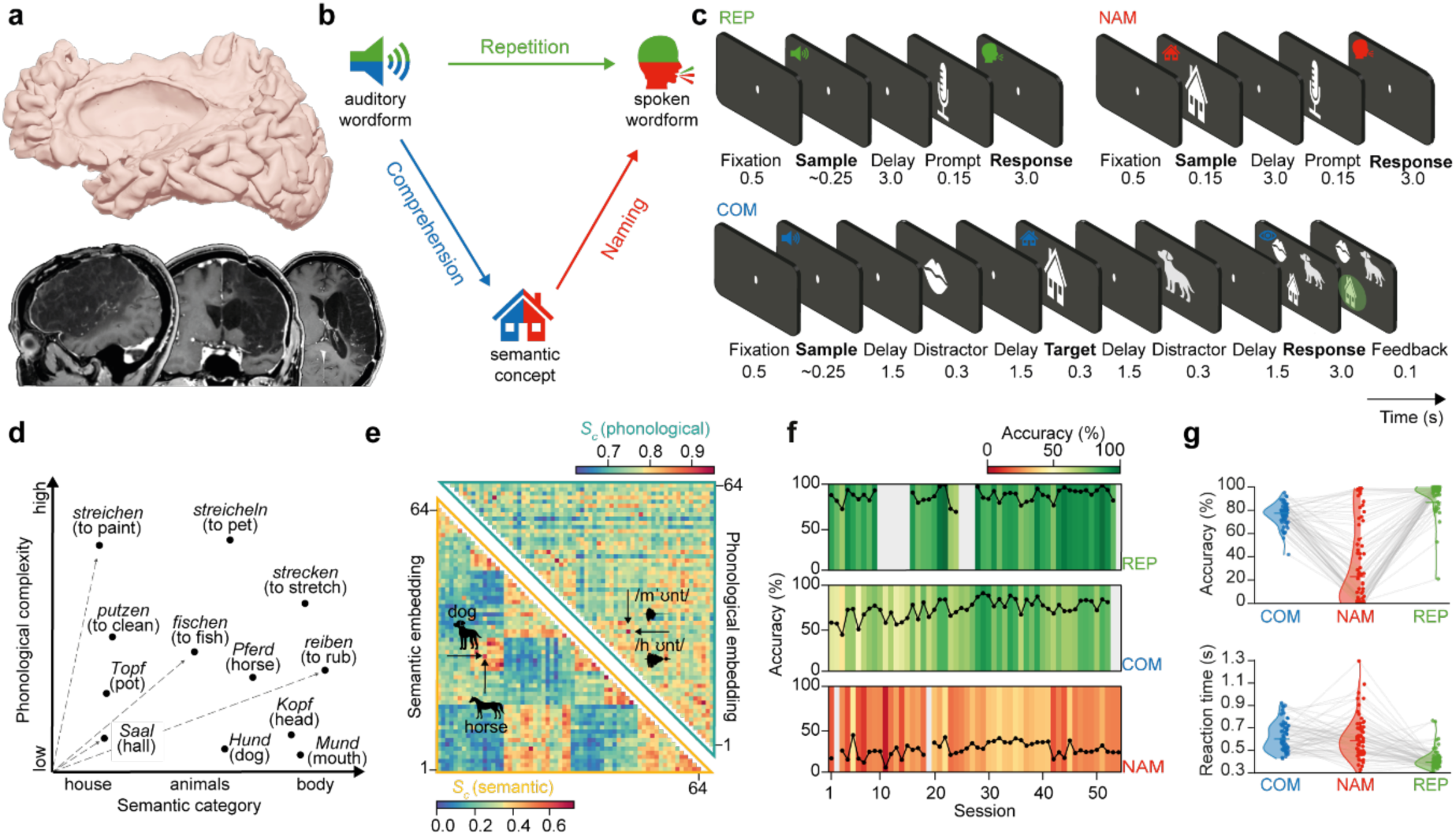
Language task design and behavioral performance. **a**, Top: 3D reconstruction of the participant’s lesioned left hemisphere. Bottom: structural MR images (T1-weighted) showing the extent of the ischemic stroke in the middle cerebral artery territory. Radiological convention: the left hemisphere is shown on the right. **b**, Schematic of the performed language tasks. **c**, Task design for word repetition (top left), word retrieval or naming (top right) and word comprehension (bottom). **d**, Schematic of the item set spanning three semantic categories (household items, animals, body parts) and different levels of phonological complexity. 12 of 64 items shown for clarity. **e**, Semantic and phonological similarity of all word pairs (bottom left and top right triangle, respectively), expressed as cosine similarity between word embeddings derived from LLMs capturing primarily semantic and phonological features (word2vec/fastText and Whisper encoder, respectively). Words are ordered by experimental subsets, and the same order is maintained in both triangles. Two example word pairs are highlighted: semantically related words (horse/dog) are similar in semantic space, while phonological minimal pairs (“Mund”/”Hund” (mouth/dog)) are similar in phonological space. **f**, Session-average accuracies for each task and recording session. **g**, Item-average accuracies (top) and reaction times (bottom) by task. Each data point represents an individual item; lines connect the same item across tasks.

We administered a set of 64 items, each drawn from one of three semantic categories (household items, animals, body parts) and associated with a German word, which varied systematically in phonological complexity (**Fig. 1d, e**; **Extended Data Fig. 1a, b; Supplementary Table 1**). Each item was represented by two visual images (different line drawings) and by two audio recordings (different speakers) of the same word, yielding a total of 128 images and 128 audio files. Items were presented in subsets of eight per recording session, with each subset used in all tasks.

Imposing distinct task demands on identical items allowed us to separate key linguistic operations such as lexical (word) retrieval in naming, phonological reproduction in repetition and semantic access in comprehension, while controlling for shared perceptual input and speech output. In line with the aphasic syndrome (**Extended Data Fig. 2**), naming was most severely impaired, while comprehension and repetition were relatively spared (27 %, 73 % and 87 % mean accuracy across sessions, respectively) (**Fig. 1f, g**). Accuracy and reaction times (RT) were only weakly correlated across tasks at the word level (Spearman’s *p*_*accuracy*_: COM–NAM = −0.32, COM–REP = 0.24, NAM–REP = −0.06; *p*_*RT*_: COM–NAM = 0.12, COM–REP = −0.20, NAM–REP = 0.08; only COM–NAM accuracy significant at p < 0.01; **Fig. 1g**), demonstrating that naming deficits were primarily driven by a selective impairment in lexical retrieval rather than downstream articulatory deficits or upstream conceptual-semantic deficits. An articulatory deficit would have resulted in comparable word-level performance in repetition and naming, a semantic deficit in comparable performance in naming and comprehension.

### Large-scale single-unit recordings

In each session, we recorded extracellular electrophysiological activity from four planar intracortical microelectrode arrays (64 electrodes each; total 256 electrodes) chronically implanted in the right-hemispheric MFG, IFG, SMG, and ANG (**Fig. 2a-c**). We determined the implantation sites based on the extent of the left-hemispheric stroke lesion and the participant’s language impairment profile. Specifically, we focused on the dorsal cortex to capture language subprocesses related to word retrieval and production^41^, the most compromised functions. Wide-band recordings were high-pass filtered (**Fig. 2d)**, spike-sorted and manually curated (**Fig. 2e, f)**, yielding well-isolated single units (SUs) and multi units (MUs) across multiple electrodes (**Fig. 2g, h)**. A total of 10,144 units were included in the analysis (MFG: 4,446 [2,226 SUs]; IFG: 3,757 [1,513 SUs]; SMG: 1,786 [811 SUs]; ANG: 155 [59 SUs]) (**Fig. 2i**). The differences in unit yield most likely resulted from differences in the extent of contact between the array and the cortical surface (determined by post-operative radiography). Because we did not track neuron identities across sessions, we consider this count as an upper bound on distinct units^42^. The fraction of electrodes with unit activity and signal-to-noise ratios (SNR) remained stable throughout the recording period (**Fig. 2j, k**).

**Fig. 2.**
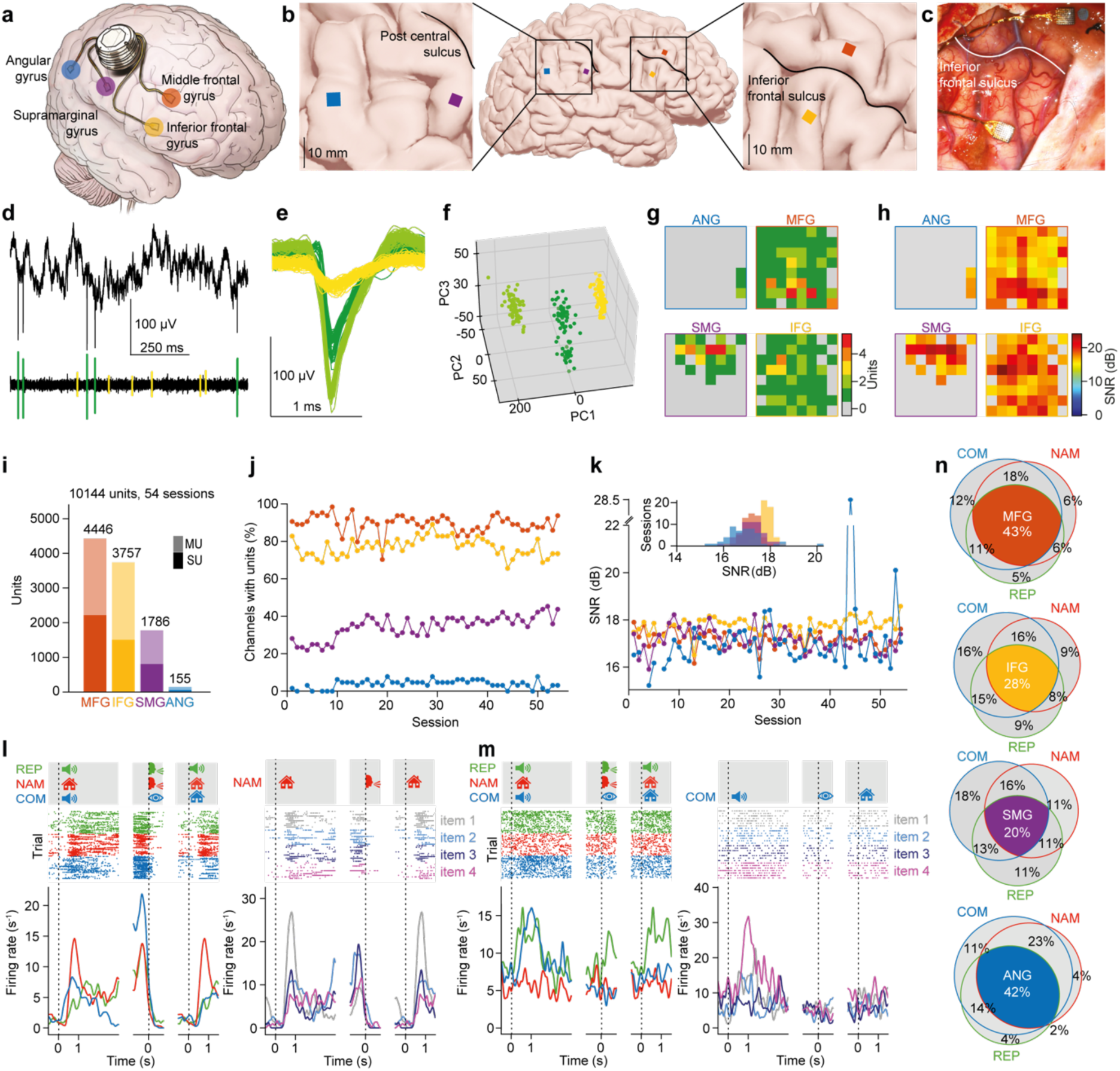
Large-scale single-unit recordings. **a**, Schematic of the microelectrode array locations in right-hemispheric prefrontal and parietal cortex. **b**, Cortical surface reconstruction and magnification of the implantation sites. **c**, Intraoperative view of the prefrontal arrays. **d**, Extracellular voltage trace of an example electrode. Top: raw signal. Bottom: high-pass-filtered spiking activity. Individual spikes are highlighted. **e**, Spike waveforms of three well-isolated units recorded on the example electrode in (**d**). **f**, Clustering of spike waveforms in principal component (PC) space. **g**, **h** Unit yield (**g**) and mean signal-to-noise ratio (SNR, averaged across all units at a given electrode; **h**) in a representative session. **i**, Unit yield per array, assuming complete turn-over from session to session. **j**, **k**, Percentage of electrodes with unit activity (**j**) and mean SNR (**k**) per array and recording session. **l**, Spiking activity of an example unit in MFG. Correct trials are sorted by language task (REP, repetition; NAM, naming; COM, comprehension; left panels) and by item in naming (right panel). Items from top to bottom are Saal (hall), Wal (whale), Reh (deer) and melken (to milk). From left to right, individual trial epochs are aligned to sample presentation, to response onset (speech onset in naming and repetition, fixation in comprehension) and to target presentation (identical to the auditory sample in repetition and visual sample in naming, but corresponding to the visual test image matching the auditory sample in comprehension). **m,** Same as (**l**) for an example unit in IFG. Right panel shows trials sorted by item in comprehension. Items from top to bottom are Tisch (table), Fisch (fish), Schaf (sheep) and scheren (to shear). **n,** Distribution of task-modulated units per array and task.

Unit activity tracked individual trial events. Firing rates were modulated both by the performed language task and by the processed items (**Fig. 2l, m** left and right panel, respectively). To determine the extent of right-hemispheric neuronal engagement in language processing, we compared baseline firing rates with trial activity. In all regions, the large majority of units were modulated by at least one language task (94 %, 87 %, 80 % and 92 % in MFG, IFG, SMG and ANG, respectively), where task modulation was defined as a significant difference in firing rate between the pre-trial baseline (500 ms epoch before sample onset) and at least two consecutive trial windows (150 ms bin width, 75 ms step, paired t-test, p < 0.05). MFG contained the largest proportion of units engaged in all three tasks (43 %), compared with ANG (42 %), IFG (28 %) and SMG (20 %) (**Fig. 2n**).

### Language-task related spiking activity

Each recorded region revealed a distinct language-task related activity profile. MFG units were strongly driven by naming and comprehension, while repetition evoked less pronounced activity (**Fig. 3a**, top). IFG units deactivated after sample onset in all tasks, then gradually recovered to baseline before verbal responses (speech onset; naming and repetition), but not nonverbal responses (eye movements; comprehension) (**Fig. 3a**, second to top). SMG activity was dominated by verbal production tasks (naming and repetition), with prominent responses to visual stimuli (**Fig. 3a**, second to bottom). ANG showed stronger modulations during comprehension than verbal production tasks (**Fig. 3a**, bottom).

**Fig. 3.**
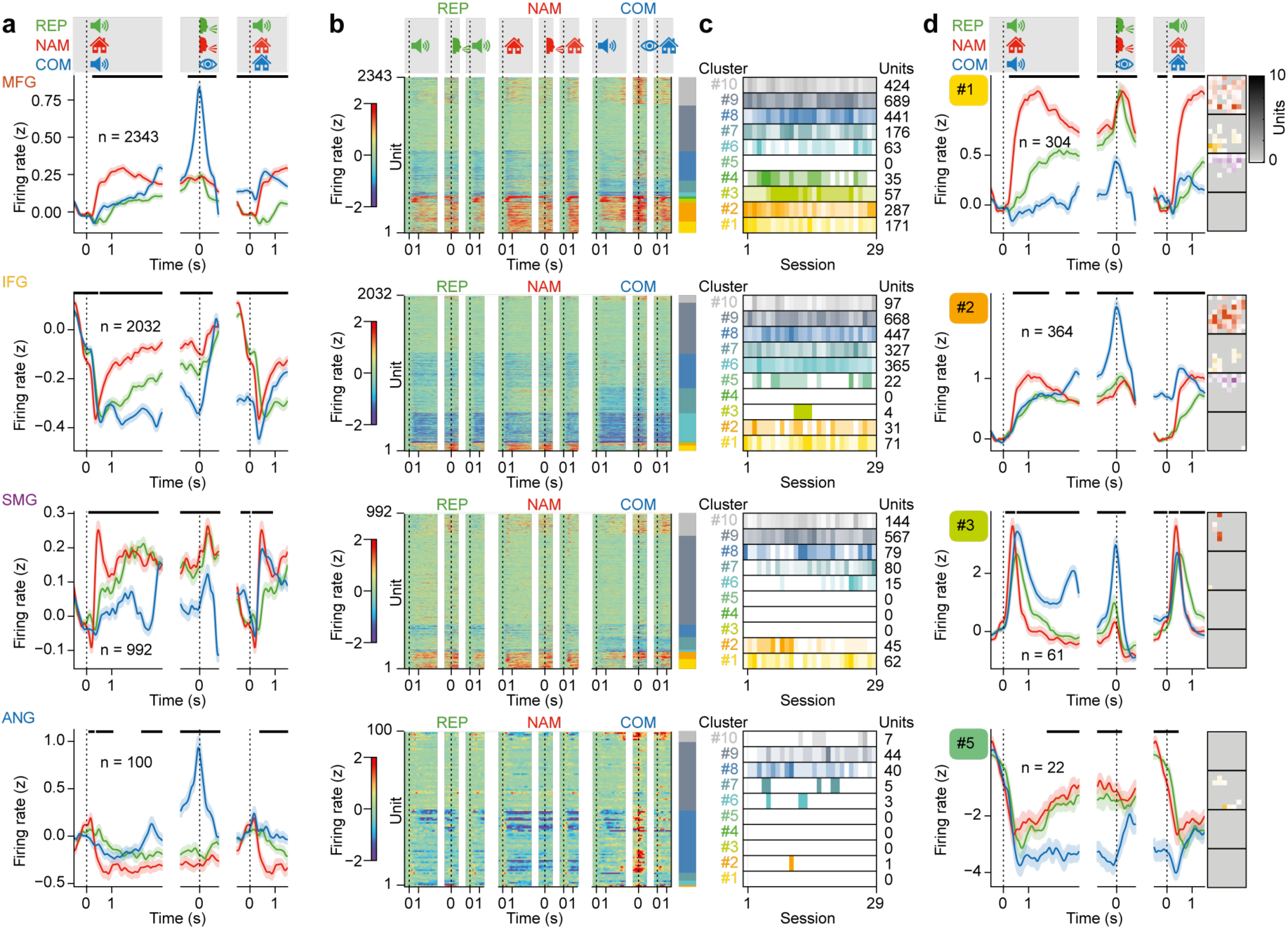
Language-task related spiking activity. **a**, Unit-averaged activity, split by region and task. Horizontal bars indicate periods with significant differences between tasks (ANOVA, p < 0.01). Error bands, SEM across units. Analyses were restricted to sessions in which all three language tasks were completed. Trials were randomly subsampled and balanced to minimally seven correct trials per item for minimally three items. Units with zero variance in baseline firing rate were excluded. Trial epochs and alignments as in (Fig. 2l, m). **b**, Hierarchical clustering of spiking activity, performed across regions, using all units from (**a**). Sorting is maintained across tasks. Colour bar on the right indicates cluster identity. **c,** Distribution of units in each cluster across recording sessions with all language tasks completed. **d,** Example clusters illustrating the diversity of response profiles. Left panels: same layout as in (**a**). Right column: spatial distribution of the units in each cluster across the four arrays. Colour saturation indicates the number of units per electrode.

In all regions, neuronal activity was significantly correlated between repetition and naming, i.e., the verbal production tasks (Pearson’s r = 0.28, r = 0.18, r = 0.17 and r = 0.24 for MFG, IFG, SMG and ANG, respectively; **Extended Data Fig. 3**). Activity in MFG and IFG was also significantly correlated between repetition and comprehension and between naming and comprehension, in line with prefrontal domain-general cognitive control. These correlations were not prominent (SMG) or absent (ANG) in the parietal regions.

To determine whether the region-specific activity reflected individual neuronal responses or instead arose from averaging across heterogeneous activity profiles, we performed hierarchical clustering of spiking activity across all units (**Fig. 3b**). Ten clusters optimally captured the principal response patterns, while remaining parsimonious and readily interpretable. Activity profiles segregated by task (often separating verbal production from comprehension; e.g., clusters 1, 5, 7, 9) and temporal dynamics (sharp event-locked peaks versus ramping or sustained responses; e.g., clusters 3, 4 versus clusters 7, 9) (**Extended Data Fig. 4**). Two notable organisational principles emerged. First, most clusters were present on multiple arrays. Only two clusters were found exclusively in MFG and IFG (cluster 4 and 5, respectively). Second, neuronal deactivation (i.e., reduction of firing rates below baseline activity) was a prominent feature and observed across all arrays, in particular in IFG. We sampled from all clusters repeatedly across the ten-month recording period, indicating they reflected stable properties of the right-hemispheric neuronal networks they were part of (**Fig. 3c**).

Four example clusters illustrate the diversity in activity profiles (**Fig. 3d**). Cluster 1 (304 units) was distributed across regions and selective for verbal production tasks. Units strongly activated following visual sample presentation in naming, but less after identical visual input (target presentation) in comprehension. Cluster 2 (364 units) was equally distributed and task-selective, but peaked primarily during the response epoch in comprehension. Cluster 3 (61 units), in contrast, showed sharp, event-locked modulation with strong regional selectivity: units in this cluster were largely confined to MFG. Finally, cluster 5 (22 units) was specific for IFG and deactivated after sample presentation. Firing rates recovered prior to speech onset in naming and repetition, but remained suppressed in comprehension.

The majority of clusters were uniformly distributed across the cortical patch covered by each array (up to 10 mm²) without evident spatial concentration (MFG: 9/9, IFG: 5/9, SMG: 6/7, ANG: 6/6; **Extended Data Fig. 4b**).

### Mixed-selective item encoding

After establishing widespread language-task related activity across the recorded regions, we asked whether individual neurons encoded specific information about the processed items, i.e., showed item selectivity. We quantified how much variance in each unit’s firing rate (expressed as explained variance ω²) could be attributed to item identity, collapsing across the two visual instances (line drawings) or the two auditory instances (audio recordings) of each item.

Populations of units were tuned to individual items in all regions, most prominently in the naming task (**Fig. 4a**). MFG and IFG showed similar selectivity profiles. In both regions, selectivity peaked after sample and target presentation and re-emerged at time of the response (**Fig. 4a** top and second to top, respectively). SMG exhibited earlier and more pronounced selectivity peaks for visually cued items in naming and comprehension, but not for auditorily cued items in repetition and comprehension (**Fig. 4a**, second to bottom). Selectivity peaks in ANG were less prominent and noisier due to the lower unit count (**Fig. 4a**, bottom).

**Figure 4.**
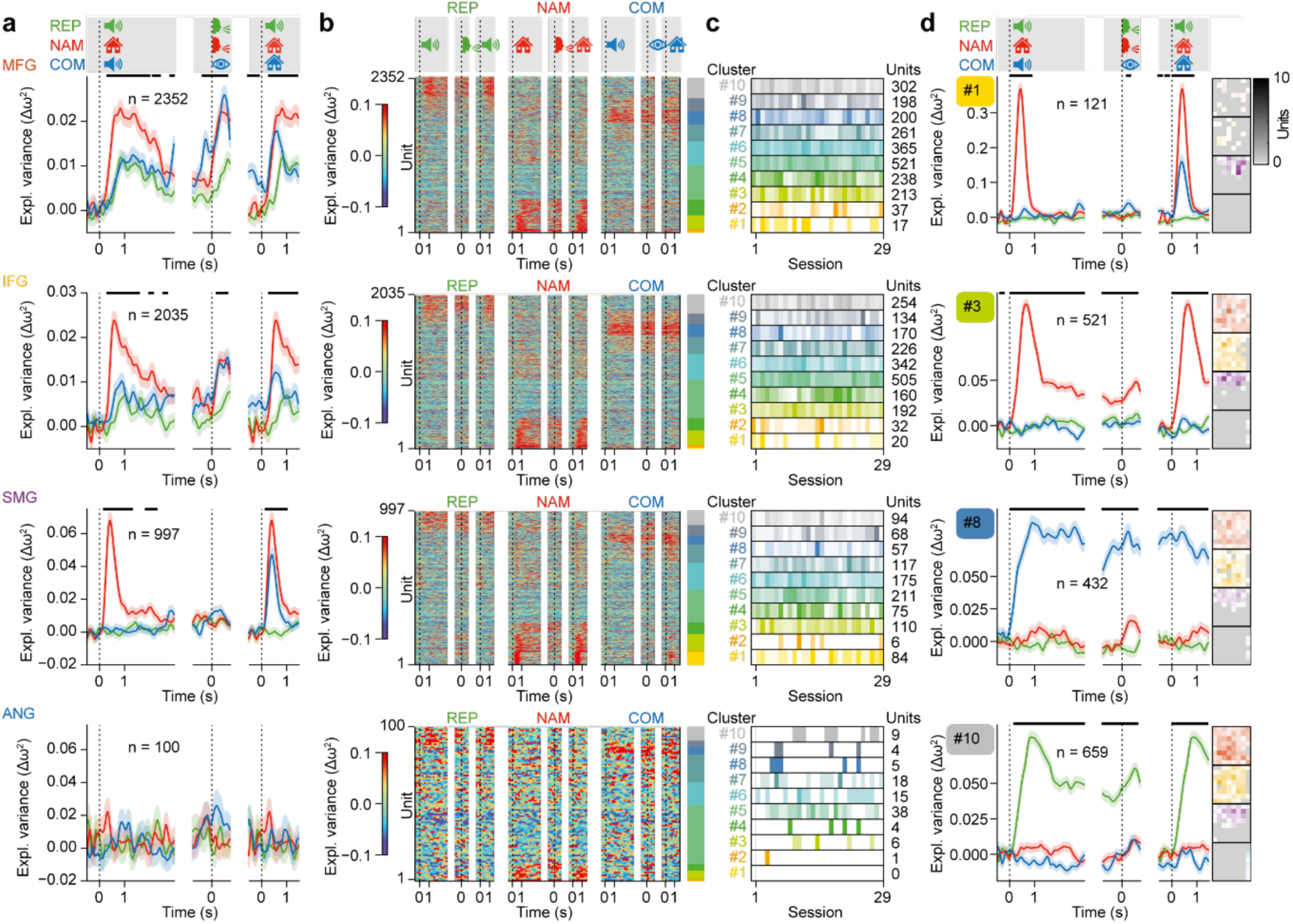
Mixed-selective item encoding. **a**, Unit-averaged item selectivity, expressed as the fraction of firing rate variance explained by item identity with shuffled control subtracted (Δω²), split by region and task. Horizontal bars indicate periods with significant differences between tasks (ANOVA, p < 0.01). Error bands, SEM across units. Differences in unit counts compared to **Fig. 3** were the result of random trial subsampling. Trial epochs and alignments as in (**Fig. 2l, m**). **b**, Hierarchical clustering of item selectivity, performed across regions, using all units from (**a**). Sorting is maintained across tasks. Colour bar on the right indicates cluster identity. **c**, Distribution of units in each cluster across recording sessions with all tasks completed. **d**, Example clusters illustrating the diversity of selectivity profiles. Left panels: same layout as in (**a**). Right column: spatial distribution of units in each cluster across the four arrays. Colour saturation indicates the number of units per electrode.

We performed hierarchical clustering of item selectivity as previously for unit activity (**Fig. 4b**, compare with **Fig. 3b)**. Strikingly, individual neurons represented item information in a highly task-specific manner. All strongly coding clusters (clusters 1, 2, 3, 4, 8, 10) carried item information in only a single task, remaining unselective for the same item in the other tasks (**Extended Data Fig. 5**). This pattern was stable across recording sessions (**Fig. 4c**). Four example clusters illustrate the diversity in selectivity profiles (**Fig. 4d**). Cluster 1 (121 units) and cluster 3 (521 units) encoded item information only following visual cueing, with strongest selectivity in naming (sample presentation) and significantly weaker-to-absent selectivity in comprehension (target presentation), despite matched sensory input. Cluster 8 (432 units) selectively encoded item information in the comprehension task, but not in the repetition task, despite matched sensory input. Finally, cluster 10 (659 units) showed the reversed pattern with selectivity to the item in repetition, but not in comprehension.

As in our activity clustering analyses (**Extended Data Fig. 4b**), item selectivity clusters were intermingled within arrays (MFG: 10/10, IFG: 9/10, SMG: 10/10, ANG: 9/9; **Extended Data Fig. 5b**). No item selectivity cluster was confined to a single region (**Extended Data Fig. 5b**).

In contrast to the activity profiles that were correlated across tasks in all regions (**Extended Data Fig. 3**), only SMG showed weak correlations for item selectivity between comprehension and naming, i.e., the tasks with common visual input (Pearson’s r = 0.04, **Extended Data Fig. 3**).

Together, these findings demonstrate mixed-selective item encoding during language comprehension and production.

### Representation of abstract lexical information

To investigate whether these item-specific signals also reflected abstract lexical (word) information in the semantic and/or phonological domain, or rather remained tied to particular sensory formats and linguistic tasks, we trained linear decoders (support vector machines, SVMs) on population spiking activity to quantify the extent to which item selectivity generalized across trial time and across tasks (**Fig. 5**).

**Figure 5.**
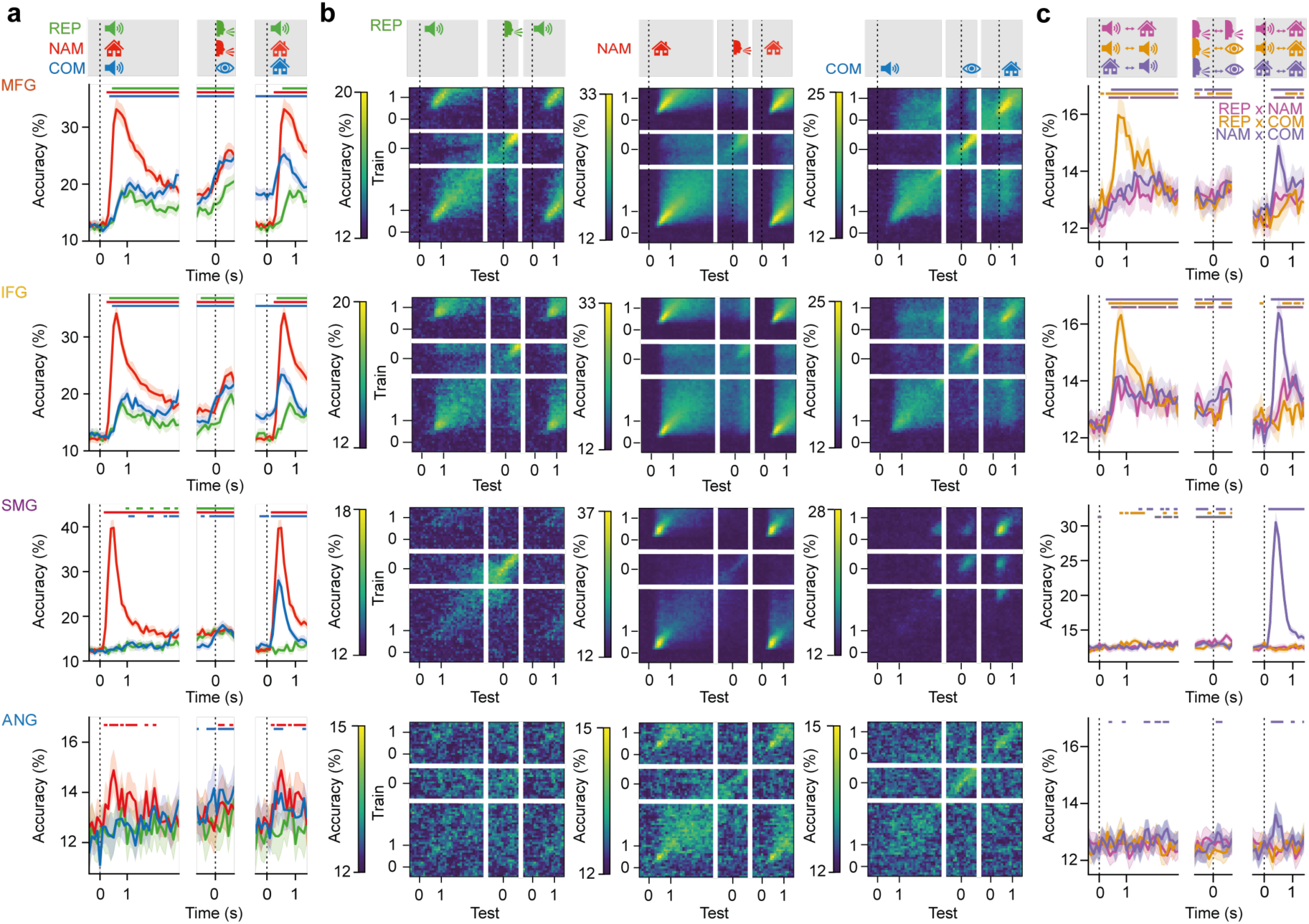
Representation of abstract lexical information. **a**, Mean within-session item decoding accuracy, split by region and task. For each session, decoding accuracy was averaged over cross-validated folds (k = 5; 80/20 split; balanced accuracy). Session-level means were then averaged across sessions. Horizontal bars indicate periods with significant group-level decoding performance (chance level 12.5 %, eight-way classification; permutation test, N=1,000; Benjamini–Hochberg FDR-controlled, q < 0.01). Error bands, 95 % confidence intervals. Trial epochs and alignments as in (**Fig. 2l, m**). **b**, Cross-temporal mean within-session item decoding accuracy, split by task (columns) and region (rows). The y-axis denotes the training window, and the x-axis denotes the testing window. **c**, Cross-task mean within session item decoding accuracy (averaged across both directions, e.g., training on repetition and testing on naming, and vice versa), split by region. Horizontal bars indicate significant cross-task generalization at the group level (12.5 %, eight-way classification; q < 0.01, corrected as in (**a**)). Error bands, 95 % confidence intervals.

In all regions and tasks, item information could be decoded above chance level when training and testing were temporally aligned (**Fig. 5a**). MFG and IFG showed similar decoding profiles (**Fig. 5a**, top and second to top, respectively). In both regions, accuracy peaked shortly after sample presentation and again around response onset. Accuracy was highest in naming, in line with the prominent item selectivity profiles observed in this task (**Fig. 4a**). Item information was also maintained during the delay epoch prior to target presentation in comprehension, despite interference by additional stimuli (**Fig. 1c**, distractor images). Repetition produced the weakest decoding, with only modest peaks after sample onset and following the response. In SMG, decoding during naming was strong (**Fig. 5a**, second to bottom). Unlike in prefrontal regions, we found no decoding following auditory sample presentation in either comprehension or repetition. Decoding in ANG was noisier due to a lower unit count but followed a similar time course as in the other regions (**Fig. 5a**, bottom).

In MFG and IFG, decoding also generalized across time with item information stably represented throughout the delay epochs (**Fig. 5b**, top and second to top, respectively). SMG showed the highest temporal specificity: decoding was tightly locked to visual stimulus presentation in naming and comprehension and to the fixation response in comprehension (**Fig. 5b**, second to bottom). In ANG, the cross-temporal structure also differed between tasks (**Fig. 5b**, bottom).

A sharp dissociation between prefrontal and parietal regions emerged in cross-task decoding (**Fig. 5c**). As expected, the strongest transfer occurred between epochs of different tasks with matched sensory inputs (e.g., auditory sample in repetition and comprehension; visual sample in naming and visual target in comprehension). However, decoding also generalized between epochs with different presentation formats (visual versus auditory) and with different response modalities (speech production versus eye movements). In MFG and IFG, decoders trained on visual stimulus activity (sample in naming and target in comprehension) achieved significant above-chance performance when tested on auditory stimulus activity (sample in comprehension and repetition) and vice versa (**Fig. 5c**, top and second to top). Surprisingly, cross-task decoding in the response epoch between naming and repetition, which share response modality (i.e., speech production), did not outperform decoding between tasks with unmatched response modality. This suggests the decoded prefrontal signals were not driven primarily by articulatory planning or vocal motor output, but instead carried semantic information about the processed items. Parietal regions showed a different pattern. SMG did not exhibit cross-modality generalization: decoders failed to transfer between visual and auditory stimulus activity (**Fig. 5c**, second to bottom). Within-modality transfer was robust, as expected, with strong decoding between visual sample in naming and visual target in comprehension. ANG showed similar modality-specificity, with cross-task decoding succeeding only when stimulus sensory formats matched (**Fig. 5c**, bottom).

In sum, these results argue that right-hemispheric prefrontal cortex also represented item information in an abstract lexical manner, beyond the modality- and task-specific formats found previously (**Fig. 4**).

### Phonological and semantic feature encoding

Next, we sought to quantify the relative contributions of phonological and semantic features to neuronal item tuning (**Fig. 4** and **Fig. 5**) at each time point in the three tasks. For this, we modelled firing rates using vectorial representations (embeddings) derived from pretrained LLMs for each word associated with the presented item (**Fig. 1e**) and quantified the extent to which phonological and semantic features explained unit activity across tasks and regions (expressed as Pearson correlation between predicted and observed firing rates^43^; **Extended Data Fig. 6**).

Word embeddings successfully predicted spiking activity. The resulting correlation traces closely mirrored the previous selectivity and decoding results (**Fig. 6a**, compare to **Fig. 4a** and **Fig. 5a**). Encoding of both semantic and phonological features was strongest in naming and weakest in repetition. We quantified individual units’ preferences for semantic or phonological features using a modulation index. In MFG and IFG, semantic coding in the target epoch dominated over phonological coding in all three tasks (**Fig. 6b, c**, top and second to top). Notably, the level of semantic dominance scaled according to the relative semantic demands of the individual tasks (strongest in comprehension, weakest in repetition). SMG, in contrast, exhibited a task-dependent reversal of feature preference. Phonological representations dominated in item naming, whereas semantic representations dominated during comprehension of the same item (**Fig. 6b, c**, second to bottom). ANG showed weaker and less consistent feature preferences (**Fig. 6b, c**, bottom).

**Figure 6.**
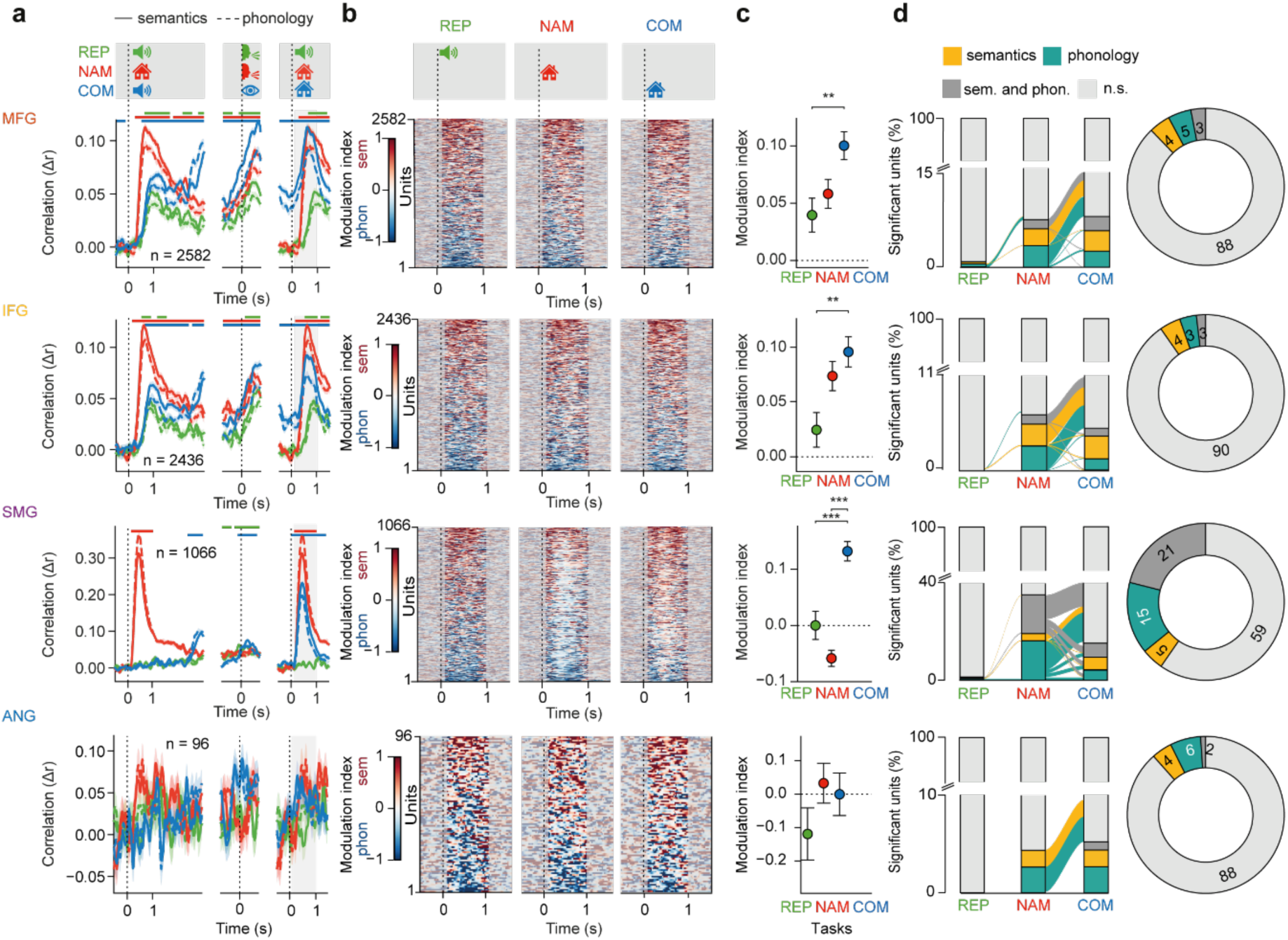
Phonological and semantic feature encoding. **a**, Time-resolved correlation traces (Pearson’s r) between model-predicted and observed spiking rates, averaged across units, for each task and region, separated by phonological and semantic model. Horizontal bars mark significant differences between phonological and semantic encoding (FDR-corrected at p < 0.01). Error bands, SEM across units. Differences in unit counts compared to **Fig. 3** and **Fig. 4** were the result of distinct trial inclusion criteria. Analyses were restricted to units co-recorded across tasks. Trials were balanced to minimally six trials per item, irrespective of accuracy. Units with non-finite correlation traces were excluded. Trial epochs and alignments as in (**Fig. 2l, m**). **b**, Unit-wise time-resolved modulation index (normalized difference between phonological and semantic model correlation strength) in the target epoch, split by task and region. Units are sorted by their average modulation index within the target epoch of the respective task. Positive and negative values indicate stronger encoding of semantic and phonological features, respectively. **c**, Task-wise modulation index in the target epoch, averaged across units. Error bars, SEM across units. Significance brackets indicate Benjamini–Hochberg FDR-corrected pairwise task contrasts from a linear mixed-effects model with a random intercept per unit (α = 0.05). * q < 0.05; ** q < 0.01; *** q < 0.001. **d**, Left: percentage of units significantly modelled (permutation test, p < 0.05) by semantic or phonological embeddings in the target epoch of the three tasks. Transitions across tasks are marked by ribbons. Only units recorded in all three tasks were included. Right: Pie charts show the percentage of significant units by region, pooled across tasks.

At the unit level, encoding of either semantic or phonological features was largely absent in repetition (max. 0.3 % significant units in all regions, permutation test, p < 0.05; **Fig. 6d**). Across naming and comprehension, SMG showed the largest proportion of language-related units (12 %, 10 %, 41 % and 12 % in MFG, IFG, SMG and ANG, respectively). Phonological features modulated the activity of most significant units in SMG, whereas semantic and phonological preference was more evenly distributed across units in the other areas (67 %, 60 %, 88 % and 67 % phonology-modulated units in MFG, IFG, SMG and ANG, respectively). These subsets were strongly task-specific, mirroring our previous selectivity analyses, indicating that naming and comprehension engage largely non-overlapping language-selective subpopulations (see also **Fig. 4b, d**).

Thus, prefrontal regions maintained a semantic preference across language tasks, while SMG flexibly reconfigured linguistic feature encoding in alignment with specific task demands.

### Population-level dynamics and representational similarity

So far, our results suggested that individual neurons in the right-hemispheric frontoparietal association cortex encoded semantic and phonological features in a task- and region-dependent manner. To determine how this information was reflected in the joint activity of the recorded neuronal ensemble, we quantified the relative contributions of task timing and item identity to population variance and tested whether population states tracked phonological and semantic relationships among words.

We used demixed principal component analysis (dPCA)^44^ to partition firing rate variance into marginalizations aligned with task variables of interest: time-dependent components (t) capturing dynamics shared across items (e.g., attention, motor preparation), stimulus-dependent components (s) capturing variance explained by differences between items, and stimulus-time interaction components (st) capturing item-specific dynamics evolving over time (**Fig. 7a**; **Extended Data Fig. 7a, b**). High time-dependent variance indicates that population dynamics are driven primarily by task temporal structure, relatively independent of item identity. High stimulus-dependent variance indicates stable item representations maintained across time. High interaction variance indicates that item representations vary with task epochs. Because dPCA performance degrades with low unit counts, ANG was excluded from these analyses.

**Figure 7.**
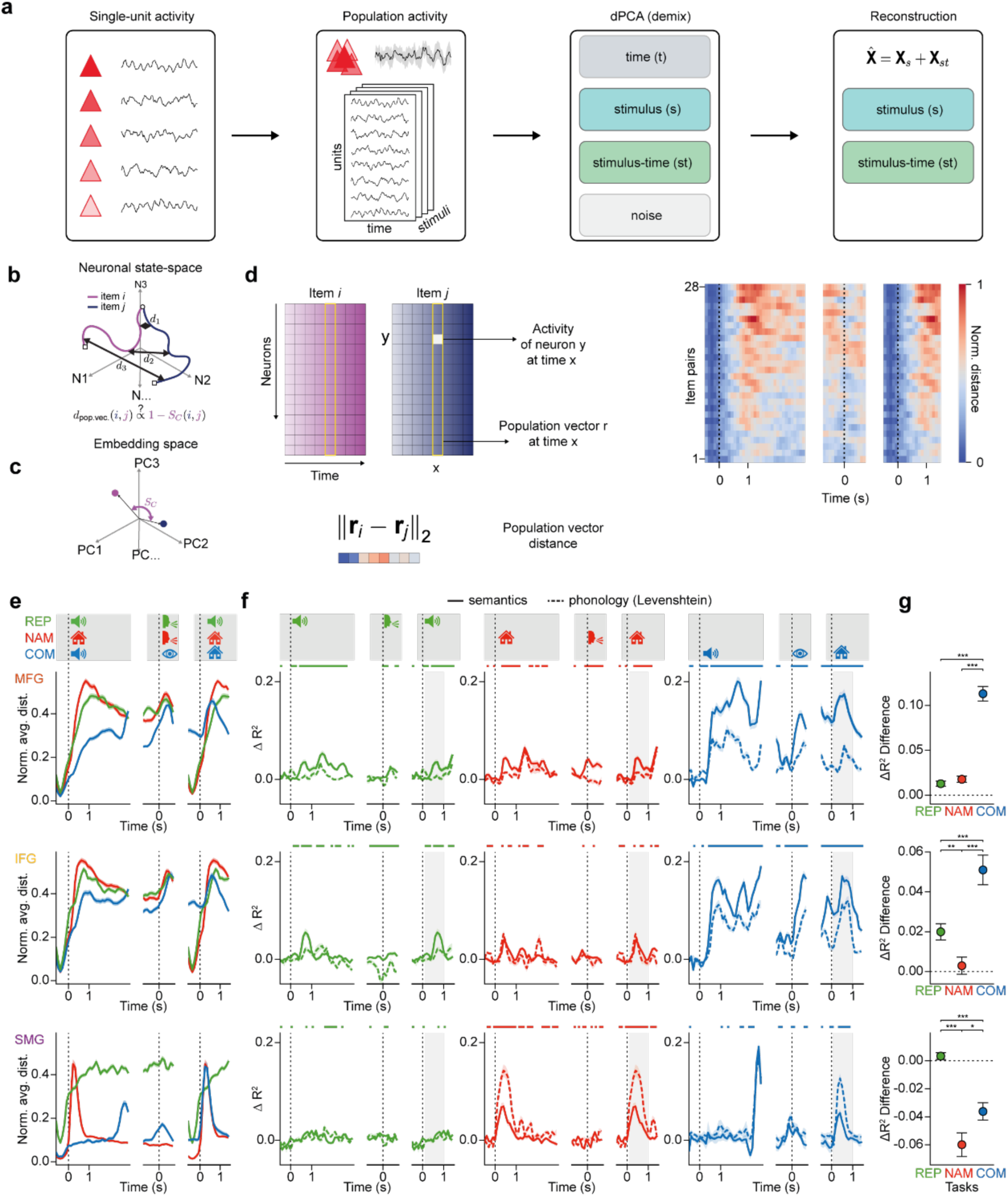
Population-level dynamics and representational similarity. **a**, Population activity X(i,t,s) was demixed with dPCA into time (t), stimulus (s), stimulus-time (st), and residual (noise) components. Activity was then reconstructed from the s and st components and represented as the trial mean. **b**, Schematic of trajectories of reconstructed population activity in high-dimensional neuronal state-space for two items, showing divergent dynamics over time. **c**, Representational similarity analysis: population vector distances between word pairs (d1, d2 and d3 in (**b**)) were correlated with the cosine distance between their corresponding embedding vectors in higher-order PC space. **d**, Left: population-vector distance. At each time point, the vector of mean firing across units for item i or j defines a point in state space. The Euclidean distance between these vectors quantifies representational dissimilarity. Right: Normalized Euclidean distances in MFG between word-pairs of one example item subset (8 words) tested in naming, sorted by mean distance, in the sample, response, and target epochs (left to right). Dashed lines mark sample, response and target onset **e**, Mean normalized population-vector distance over time, split by region and task. **f**, Time-resolved difference in test-set coefficient of determination (ΔR²) between GLMs predicting population vector distances and shuffled-label controls, split by task and region, for semantic features (fastText embeddings, cosine similarity in 20-dimensional PCA space) and phonological features (Levenshtein distance). Error bands, SEM across cross-validated folds (n = 100; 80/20 split). Horizontal bars indicate significant differences between semantic and phonological ΔR² curves per time bin (two-sided paired t-tests, FDR-corrected q < 0.05). Grey boxes indicate the target epoch of interest used in (**g**). **g**, Mean ± SEM of ΔR² difference (fastText minus Levenshtein) averaged over the target epoch (0.1 – 1.0 s post-alignment), computed across cross-validated folds. Significance brackets indicate pairwise comparisons between tasks (Welch t-tests, FDR-corrected q-values). * q < 0.05; ** q < 0.01; *** q < 0.001.

In MFG and IFG, time components explained the largest proportion of variance (69 % and 52 % across tasks, respectively; **Extended Data Fig. 7c-e**, top and middle), reflecting their role in general task timing and structuring. In contrast, interaction components accounted for nearly half of the variance in SMG (47 %, 52 % and 48 % in repetition, naming and comprehension, respectively), indicating that item-related responses were tightly coupled to specific task epochs rather than expressed as temporally stable representations (**Extended Data Fig. 7c-e,** bottom). Across all regions, naming produced the largest item-related variance (s + st combined: 34 %, 47 % and 71 % in MFG, IFG, and SMG, respectively), consistent with the stronger selectivity and decoding observed previously in this task (**Fig. 4** and **Fig. 5**).

Reconstructing population activity using only stimulus-related components (s + st) yielded stimulus-specific trajectories in neuronal state space (**Fig. 7b**). In all regions, trajectories for different items diverged during both sample and response epochs. We quantified this coding dissimilarity by computing Euclidean distances between population vectors at matched time points across all item pairs (**Fig. 7c**). To determine whether these population-level dissimilarities reflected phonological or semantic structure, we performed representational similarity analysis (RSA), testing whether word-pair differences in embedding space predicted population-vector distances (**Fig. 7d**).

Representational dissimilarity increased after sample onset and remained elevated throughout the trial. MFG and IFG showed similar time courses with stable, persistent coding across the delay epochs, whereas population dynamics were more trial-epoch specific in SMG (**Fig. 7e**).

Across all regions and tasks, differences in both phonological (Levenshtein distance) and semantic (fastText) features predicted out-of-sample population distances above chance (**Fig. 7f**; **Extended Data Fig. 7f).** Encoding was stimulus-locked across regions, strongest in comprehension, and minimal in repetition. In MFG and IFG, RSA revealed both sample- and response-aligned peaks, with comprehension showing the most sustained encoding across both epochs. In SMG, encoding was tightly locked to visual stimulus onset with little response-aligned structure. Feature dominance differed by region: in MFG and IFG, semantic features dominated across tasks, whereas in SMG, phonological features contributed more strongly in both naming and comprehension (**Fig. 7g**).

Together, these results show that task timing and item identity contributed separable sources of variance to coordinated population activity. SMG networks emphasized stimulus-by-time interactions, with item representations tightly bound to specific task epochs and dominated by phonological structure. Prefrontal networks (MFG and IFG), in contrast, maintained more temporally independent, semantically weighted representations alongside response-aligned dynamics.

## Discussion

This work constitutes the first systematic investigation into the single-neuron correlates of language functions in the non-dominant right hemisphere of the human brain. Comprehension, retrieval and articulation of words widely engaged the recorded regions. Lexical, word-level information was readily decodable from both single-neuron activity and population-level dynamics, with semantic features preferentially accounting for prefrontal activity and phonological features for parietal activity. The representational format of lexical information was mixed-selective: individual neurons functionally clustered into distinct subpopulations that encoded language items in a highly task-specific manner, despite correlated firing patterns across tasks. Together, our results suggest that right-hemispheric neuronal populations are involved in sensory-to-semantic and semantic-to-articulatory transformations that are at the core of language processing.

Only little more is known about the single-neuronal elements of language in the dominant left hemisphere. Nevertheless, several of our findings relate to and extend results from recent intracranial studies in the homotopic regions of the left brain.

In left MFG, single neurons were found to encode the planned phonetic composition and sequential structure of utterances before articulation^8^ as well as the semantic content of words during sentence comprehension^3^. In our recordings, activity in right MFG similarly reflected phonological and semantic word features, suggesting that accessing lexical representations during word retrieval and comprehension also engages the non-dominant hemisphere. Notably, we observed stronger item decoding in naming than in repetition in all recorded regions, arguing that semantic structure is also the primary driver of right-hemispheric responses in other prefrontal regions and in parietal cortex.

Single-unit recordings in epilepsy patients have recently identified neurons in the right IFG with comparable responses to spoken and written words^45^. We similarly found correlated activity across tasks. Selectivity for individual items, however, was clearly task-specific, consistent with the prefrontal cortex’s role in goal-oriented behaviour and flexibly representing information according to current task demands. Interestingly, suppression, rather than elevation, of spiking activity following sample presentation was a dominant feature in right IFG, a phenomenon that parallels recent electrocorticographic recordings of neuronal population activity during speech planning, predominantly in the right hemisphere^45^. This inhibition could be unrelated to language per se and instead reflect the suppression of irrelevant inputs into task-specific networks, in the sense of inhibitory control. Rather than contradictory, the activity-selectivity contrast suggests that the right prefrontal cortex contributes not only to representational content but also to modulating the neural landscape in which language processing unfolds.

In the left parietal cortex, single SMG neurons of individuals with tetraplegia have been found to encode phonetic-phonological word structure across auditory comprehension, reading, internal speech and vocalized speech^9^. We observed the same in right-hemispheric parietal spiking activity. SMG activity was more phasic and modality-sensitive than in prefrontal cortex. SMG also contained the highest proportion of units showing significant modulation by phonological features. Unlike frontal regions, SMG neurons prioritized phonological processing in naming and semantic processing in comprehension, suggesting stronger integration of language sounds and language meaning in parietal cortex.

Our work speaks directly to longstanding questions about the nature and specificity of right-hemisphere recruitment in chronic aphasia following left-sided brain damage, especially stroke. At the level of individual neurons, we demonstrate here that the right hemisphere can express structured, task-dependent linguistic representations, rather than diffuse, non-specific activation. These findings support the view that the non-dominant hemisphere is linguistically responsive and exhibits functional organisation and regional differentiation for language^46^, with considerable symmetry to the left, in the sense of a language “shadow”^47,48^.

Nevertheless, the distributed organisation we observed with many features shared across prefrontal and parietal cortex suggests that right-hemispheric language-sensitive regions are less specialized than their left-hemisphere counterparts, a property that may confer flexibility in the context of reorganisation after brain damage. Consistent single-unit responses over ten months indicate that this representational structure is a stable feature of the chronic state. However, to what extent right-hemispheric resources can be leveraged to drive behavioral gains for patients with aphasia likely depends on how effectively language-responsive units functionally integrate into the reorganised network^49^. This question has direct implications for both language rehabilitation and neurotechnological approaches targeting these networks. Future work will be essential for understanding what determines whether local representational structure can be translated into measurable recovery.

For example, recent electrocorticographic recordings have demonstrated clustering of language activity on the sub-centimeter scale^50,51^, which we now resolve at the level of single neurons. Activated and suppressed units were intermingled within individual arrays, with distinct subpopulations selectively recruited depending on task demands. This fine-scale structure has implications for intervention design and calls for mechanistically specific approaches. Neurofeedback, closed-loop neuromodulation and decoding, selectively engaging neuronal subsets that carry linguistic information while preserving the domain-general (inhibitory) dynamics, could significantly augment residual language capacities in chronic aphasia beyond what behavioral therapy alone currently can achieve.

We cannot tell whether the language-task activity we observed reflects post-stroke reorganisation, the expression of premorbid bilateral organisation, or both. More work is necessary to determine how right-hemispheric representational (pre-)structure relates to the trajectory of language recovery, a question for which our characterization of single-neuron language coding in the chronic, stationary phase of aphasia provides a useful starting point.

Also, as we report data from a single participant, the extent to which our observations generalize remains to be established. However, single-case studies address a different explanatory level than group-based designs. Aphasia is a heterogeneous clinical syndrome with mechanisms varying between individuals. In BCI development, inference at the level of the single subject will therefore be essential, as decoding must ultimately be optimized within a single brain. An *n* = 1 design enables deep phenotyping at spatial and functional resolutions inaccessible to non-invasive neuroimaging, linking behavior and neuronal dynamics within the same system rather than across group averages. Thus, the results presented here do not constitute a theory of aphasia in general, but a mechanistic account with far-reaching translational relevance.

## Methods

### Participant

The participant was a 51-year-old female with higher education who had suffered an ischemic stroke in the left middle cerebral artery territory (M1 occlusion) six years prior to study enrollment. The stroke lesion comprised extensive necrosis of the frontal, temporal, and parietal cortices, resulting in near-complete destruction of the left perisylvian language network. The acute post-stroke phase was marked by global aphasia and apraxia of speech. At study onset, standardized clinical assessments, including the Aachener Aphasie Test (AAT^52^) and Aphasia Checklist (ACL^53^) indicated non-fluent (“Broca’s”) aphasia and moderate apraxia of speech (Hierarchische Wortlisten^54^). Expressive language was severely impaired; auditory comprehension was preserved for single words and simple syntax but declined with increasing syntactic complexity; written language deficits paralleled spoken impairment, with absent functional word-writing. Severe lexical retrieval deficits were confirmed by the Bielefelder Wortfindungsscreening^55^. Non-verbal cognition, including non-verbal semantic processing (NVST^56^), and orofacial praxis were intact. Longitudinal monitoring indicated that the language profile was stable (**Extended Data Fig. 2**). The participant received ongoing speech therapy targeting apraxia of speech during the study, independent of experimental tasks.

All procedures were conducted in accordance with the Declaration of Helsinki and were approved by the ethics committee of the TUM School of Medicine and Health (2018-489-S-KK). The participant provided written informed consent after the nature and potential risks of the study had been explained in accessible language. Consent was reaffirmed repeatedly throughout the study. Participation was voluntary, and the participant could withdraw at any time without consequences for clinical care.

### Electrode placement

Implantation sites were planned based on presurgical mapping: fMRI and DTI together identified preserved structural and functional connectivity within the right hemisphere, including coupling between the inferior frontal and inferior parietal regions. Transcranial magnetic stimulation (TMS) using a word–picture mapping paradigm did not yield conclusive localization evidence, as stimulation at any cortical site produced performance decrements. These findings collectively indicated that residual language function was likely sustained by a broadly distributed right-hemispheric homotopic network, and guided selection of implantation targets.

Four 64-electrode, 1.5-mm-long silicon planar microelectrode arrays (“Utah arrays”, Blackrock Neurotech) connecting to a single pedestal were chronically implanted into the right middle frontal gyrus (MFG), inferior frontal gyrus (IFG), supramarginal gyrus (SMG) and angular gyrus (ANG) under general anesthesia. In MNI space, the array centroids are located at 47.75, 31.12, 34.54 (MFG; border of BA6 and BA8, Allen Human Brain Atlas^57^), 61.66, 18.12, 18.50 (IFG, BA44, 80 % probability, Jülich Brain Atlas^58^), 66.78, –24.36, 30.74 (SMG, BA40, 69 % probability), and 62.18, –49.67, 37.90 (ANG, BA39, 68 % probability).

### Task design and stimulus material

Three single-word-level language tasks were designed to dissociate cognitive subprocesses of language production and comprehension while accommodating the participant’s language abilities. All tasks were administered with eye-tracking–based control: trials advanced only upon continuous fixation of a central fixation point, ensuring stable visual attention prior to stimulus onset. Tasks were blocked and run in pseudo-random order within the recording session.

#### Naming task

Each trial was initiated by a 500-ms fixation period. Upon successful fixation acquisition, a sample image was presented centrally for 150 ms, followed by a 3,000-ms delay. A microphone icon (150 ms) served as the response prompt, at which point fixation was released and participants had 3,000 ms to vocalize the target word prior to inter-trial interval onset.

#### Repetition task

Trial structure was identical to the naming task, with the exception that an auditory sample, i.e., a pre-recorded spoken word (mean duration: 250 ms) delivered binaurally via headphones, replaced the visual sample.

#### Comprehension task

Each trial was initiated by a 500-ms fixation period, followed by auditory sample presentation. Participants maintained fixation throughout a subsequent 1,500-ms delay, after which three images were presented sequentially (150 ms each; 1,500-ms inter-stimulus intervals) in randomized order, comprising the target image (semantically congruent with the auditory sample), one semantically or phonologically related distractor, and one unrelated distractor drawn from the item set. During the final response epoch, all three images appeared simultaneously at equidistant positions along a circular array. Target selection was operationalized as a 500-ms fixation, with response classification (target, related error, unrelated error) determined from fixation location. Accuracy feedback was delivered audiovisually via a green or red circle overlaid on the selected image, paired with a corresponding tone.

#### Trial alignment and contrasts

Neuronal activity was analysed in three time windows aligned to key task events. The sample epoch was aligned to sample onset and extended from 500 ms before to 3,000 ms after onset of the first stimulus (–500 to +3,000 ms). The response epoch was aligned to voice onset in the naming and repetition tasks, and to fixation of the selected image in the comprehension task. The response epoch extended from 750 ms before to 750 ms after the response (–750 to +750 ms). Lastly, for the target epoch, we aligned to sample onset in the naming and repetition tasks, and to onset of the target, i.e., the image matching the auditory sample, appearing as one of three sequentially presented stimuli. The target epoch extended from 500 ms before to 1,500 ms after target onset (–500 to +1,500 ms).

#### Item set

We administered items in subsets of 8 per recording session, with each subset used across all tasks within a session. The full set comprised 64 items balanced by semantic category (household items, animals, body parts) and grammatical category (nouns and verbs) within recording subset. Each item was represented by two distinct line drawings and two spoken-word recordings from different speakers (128 images and 128 audio files in total). Within each session, each image and audio file was presented ten times per task, yielding 20 (2×10) trials per item per task and 160 trials per task per session.

### Data collection and preprocessing

#### Recording setup

The participant was seated at a desk and viewed the experimental display (Samsung LED, 23,6’’), viewing distance 57 cm) through a head-mounted eye tracker equipped with an integrated chin rest for head stabilisation. MonkeyLogic 2 (NIMH) running on a dedicated PC was used for experimental control and behavioral data acquisition. Behavioral time stamps were transmitted to the NSP system for parallel logging of neuronal data and behavioral events. Visual stimuli were presented centrally on the display. Eye position was required to remain within a fixation window of 3° radius. Auditory stimuli were delivered via headphones (Sennheiser HD 560). A microphone (Sennheiser ME 36) positioned in front of the display recorded all vocal responses. Extracellular voltage signals were sampled at 30 kHz, filtered and digitized using a 256-channel Neuroport Neural Signal Processor (NSP, Blackrock Neurotech) in conjunction with a Neuroport 256-channel headstage attached to the pedestal. The digitised signals were transmitted via digital hubs to the NSP and a recording PC for storage and offline analysis.

#### Spike sorting and unit classification

Raw signals were bandpass filtered (250 – 7,500 Hz) and common-average referenced within each array by subtracting the array mean from each channel. Spikes were sorted into unit clusters using Kilosort 1.0^59^. Spike quality was assessed by calculating the Mahalanobis distance of each spike to its cluster centroid and fitting a chi-squared distribution to these distances. Spikes falling below a threshold of 1/N (N = total spikes in the unit) were excluded as noise^60^. Units were retained if they met minimum criteria for mean firing rate (≥ 0.1 Hz) and signal-to-noise ratio (SNR ≥ 15 dB). All units were additionally inspected manually to confirm biologically plausible waveforms and to classify them as single- or multi-unit activity. Classification was based on peak-to-peak amplitude distribution and stability, inter-spike interval violations (assuming a 1-ms refractory period), and cluster separation in principal component space. Unit identity was not tracked across sessions; thus, potential turnover between sessions was not controlled. The proportion of electrodes with unit activity was computed as the number of electrodes recording at least one unit divided by the total number of electrodes on that array. The signal-to-noise ratio was calculated as

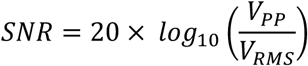

where V_pp_ is the mean peak-to-peak amplitude across spikes of a unit and V_RMS_ is the root-mean-square (RMS) voltage of the filtered signal on its electrode.

#### Behavioral scoring and response timing

Each trial was evaluated for response accuracy and response time. For naming and repetition, voice onset was estimated from the microphone trace using a custom amplitude-based detector. Voice onset was defined as the start of the first voiced segment. Voiced segments were manually transcribed and scored as correct if the transcription matched the target word exactly. For comprehension, response time was determined by the moment the eye trace first reached the logged image.

#### Cross-task behavioural covariance

To compare performance across tasks, we calculated pairwise covariation of accuracy and response times between naming, repetition, and comprehension. For each item, mean accuracy was computed across sessions separately for each task, and associations were assessed using Spearman correlation coefficients. Two-tailed significance tests were applied to both accuracy and response time measures.

### Task-modulated units

Task-modulation was defined by a significant difference in firing rates during at least one task epoch compared to baseline. Firing rates were assessed across all three alignment windows for each task, and units were classified as task-modulated if they showed significant difference to baseline for at least two consecutive 150-ms windows stepped by 75 ms (paired t-test, p < 0.05).

### Activity and selectivity profiles

#### Trial selection and preprocessing

For activity and selectivity analyses, only correct trials were included, and only sessions with at least seven correct repetitions of at least three items in all tasks were analysed. For each session, trials of items with at least seven correct repetitions in all tasks were retained, and subsampled to equalize counts per item. Lastly, units with zero baseline variance were excluded. Due to random-trial subsampling, the number of included units varies slightly across analyses.

#### Activity

Firing rates were computed in 50-ms bins and smoothed with a 50-ms Gaussian kernel. Z-scored firing rates were obtained by subtracting the trial’s baseline mean firing rate (−500 to 0 ms pre-sample) and dividing by the baseline variance across trials.

#### Selectivity

Item information was quantified using the *ω*^2^ explained variance measure. The extent to which a neuron’s firing rate varied with the items was calculated using 250-ms windows and 20-ms steps as

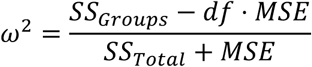

with *SS*_*Groups*_ denoting the sum-of-squares between groups (items), *SS*_*Total*_ the total sum-of-squares, df the degrees of freedom, and MSE the mean squared error. For the same timepoints as for *ω*^2^, we calculated 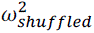 by shuffling trial labels 25 times, calculating *ω*^2^ percent explained variance and averaging across shuffles. The selectivity measure Δ*ω*^2^ was then defined as 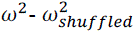.

#### Task differences

Horizontal black bars in plots of activity or selectivity profiles indicate epochs during which at least one task differed significantly from the others (150ms windows, one-way ANOVA, p < 0.01). To examine correlations between tasks, Pearson product–moment correlations were computed for all pairwise task combinations at the single-unit level. For each array and task combination, the distribution of correlation coefficients was tested against zero (one-sample t-test, p < 0.01).

#### Hierarchical clustering of activity and selectivity profiles

For each unit, z-scored PSTHs across alignment windows and tasks were concatenated to form activity profiles. Explained variance computed over the same windows was concatenated analogously to generate item selectivity profiles. Units were clustered hierarchically using Ward’s linkage and Euclidean distance, separately for activity and item selectivity profiles. To enable comparison between profile types and capture modest but potentially informative structure, the number of clusters was fixed at ten.

#### Spatial autocorrelation analysis

Spatial organisation of clusters across arrays was assessed using Moran’s I, which quantifies spatial autocorrelation (−1 = dispersion, 0 = random, +1 = clustering). For each cluster, the observed I was compared to a permutation-based null distribution generated by shuffling electrode locations 100 times (permutation test, 99^th^ percentile). To avoid confounds arising from unit turnover and spatial differences in signal quality (e.g., varying numbers of unit-recording electrodes), shuffling was performed within sessions and arrays.

### Representation of abstract lexical information

Decoding analyses were performed on windowed firing rates (200 ms window with 100 ms overlap) from all units, trials were balanced per item (minimum six trials per item, irrespective of accuracy. Several machine-learning pipelines, including linear discriminant analysis (LDA), support vector machines (SVMs) with linear and radial basis function (RBF) kernels, k-nearest neighbours (KNN), random forest (RF), and extreme gradient boosting (XGB) were evaluated. The linear SVM classifier yielded the highest and most stable decoding accuracy while maintaining computational efficiency and was therefore used for all subsequent analyses. Features were standardised using a StandardScaler to improve convergence.

#### Cross-validation and performance metrics

Firing rates were computed for each window and unit, yielding a feature vector of dimensionality 1 × n, where n is the number of units. All decoding analyses employed five-fold stratified cross-validation, in which the dataset was randomly partitioned into five subsets with four folds used for training and one for testing in each iteration, ensuring proportional class representation across folds. Performance was evaluated using balanced accuracy to account for potential class imbalances.

#### Within-session decoding

Within-session decoding established a baseline for the extent to which neuronal activity within a single session could be used to decode items directly. Each 200-ms window was decoded independently using five-fold stratified cross-validation. Accuracies were averaged across folds within sessions and then across sessions for each cortical region and task. To assess whether performance significantly exceeded chance, a hierarchical group-level permutation test was implemented. For each window, an empirical null distribution was constructed by shuffling predicted labels within each cross-validation fold over 1,000 iterations. For each permutation, a null group mean was computed by averaging first across folds and then across all sessions. Empirical p-values were calculated as the proportion of null values exceeding the observed group-mean accuracy. To account for multiple comparisons across time windows, p-values were adjusted using the Benjamini–Hochberg false discovery rate (FDR) procedure, with significance defined at an FDR-controlled threshold of q < 0.01.

#### Cross-temporal decoding

Cross-temporal decoding was performed by training the classifier on neural activity from one time window and testing on all other windows within the same session and task. Each 200-ms window served as both training and testing input in a full pairwise comparison matrix. Five-fold stratified cross-validation was applied to each training–testing combination. To prevent data leakage, trials used for training in a given window were excluded from all testing windows. For each window, decoding accuracies were averaged across folds within each session and then across sessions for each cortical region and task.

#### Cross-task decoding

For cross-task decoding the classifier was trained on one task and tested on another, using the corresponding task-aligned 200-ms windows. Only recording sessions with all tasks completed were included. Units were restricted to those stably recorded across all three tasks on the same day. Two nested five-fold stratified cross-validation procedures were used: one to define training folds in the source task and one to define testing folds in the target task, with 80 % of trials used in each case. For each window, decoding accuracies were averaged across folds within each recording day and then across all recording days for each cortical region. The mean of both training–testing directions was then computed, yielding a single cross-task generalisation metric for each task pair. Statistical significance was determined using the permutation and FDR-correction procedures described for within-session decoding.

### Phonological and semantic feature encoding

#### Multimodal word embeddings

To construct a multimodal feature space for each stimulus, we integrated two sources of lexical descriptors: Whisper encoder embeddings^61^ were used to represent the acoustic features of each word. Each audio file (two speakers per word) was processed separately. Voice onset and offset were detected using a custom amplitude-based detector (absolute waveform, 5-sample moving average; threshold = 0.05 on the normalized −1 to 1 scale), and the waveform was segmented to the detected speech interval. Segments were then padded or trimmed to a fixed duration (490 ms), resampled to 16 kHz and converted to the Whisper input format. Each segment was passed through the encoder of the pre-trained Whisper large-v3 model (bypassing the decoder) to obtain hidden states capturing acoustic content. A fixed-length embedding was created by concatenating the final 20 time steps of the last hidden state, aligned to the detected onset. These embeddings are henceforth referred to as phonological embeddings. FastText embeddings (German Common Crawl model, cc.de.300; 300-dimensional)^62^ were used to represent the distributional semantics of the words. FastText was chosen over the Whisper decoder because the decoder embeddings incorporate contextual predictions from preceding and following words, which were absent in our single-word tasks. For each word in our item set, the corresponding fastText vector was retrieved directly by look-up.

We modelled spike count vectors as a function of word embeddings using time-resolved linear regression, performed separately for each embedding type and unit^43^. Units were required to be co-recorded across tasks (i.e., in the same recording session) and trials were balanced per item (minimum six trials per item, irrespective of accuracy).

To construct two separate encoding models, we used the first 50 PCs of the fastText or the Whisper embeddings for each word. These models were applied to neuronal data during the three task alignment windows (epochs). Within each epoch, spike trains were converted to firing rates using a sliding window approach (200-ms window, 25-ms step). Spike counts were divided by window size (in seconds) to obtain firing rates and smoothed with a Gaussian kernel (σ = 1). An ordinary least-squares (OLS) linear regression model with an intercept was fitted to predict binned firing rates from the embedding features. Coefficients were estimated by OLS independently for each time bin:

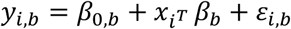

Where *i* indexes trials, *b* indexes time bins *x*_*i*_ is the embedding vector (fastText or Whisper) for trial

*i*, *β*_0,*b*_ is the intercept for bin *b* and *β*_*b*_ are the regression coefficients for bin *b* and *εi*, *b* is the residual.

For each unit and epoch, models were trained on 80 % of the trials and predicted firing rates on the held-out 20 %. A single fixed train-test split (test size = 0.2, random seed = 42) was reused across all units within the same session and epoch. Stratification was performed by item.

This procedure yielded a temporal response profile of model performance throughout the three epochs. For each time bin *b*, the vectors *y*_*true*_[:, *b*] and *y*_*pred*_[:, *b*] across test trials were extracted, and the Pearson correlation coefficient was computed as

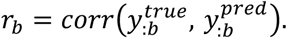

Bins in which either vector had zero variance across trials were excluded, and units with any excluded bins were removed from further analysis. The resulting correlation curves reflect within-lexeme prediction accuracy (since training and test sets included trials from the same words) and allowed dissociation of the contributions of semantic and phonological features to firing-rate modulations across task phases, rather than generalisation to unseen words.

For each unit and epoch, null distributions were generated by repeating the analysis with 1,000 random permutations of the training data under a fixed train–test split. In each permutation, the pairing between each trial’s embedding features and its neuronal response was randomly shuffled within the training set, while the held-out test trials remained fixed. Model predictions on the unpermuted test data were recomputed for each permutation, and per-bin correlations were stored to form empirical null distributions. Line plots and heatmaps display the correlation time course with the permutation mean subtracted, per model and window, averaged across units within each region (mean ± SEM)

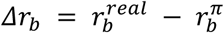

#### Region-level comparison of semantic and phonological models

To compare semantic (fastText, FT) and phonological (Whisper, WH) models over time at the region level, we performed a paired, sliding-window test on permutation-subtracted traces. For each unit, model, window, and time bin we use the permutation-subtracted correlation *Δr*_*b*_. Within each alignment window, *Δr*_*b*_ was averaged within 250-ms windows with 50 % overlap (bin width = 25 ms yielding 10-bin windows, 5-bin steps). For each task and time window, we formed two vectors across units:

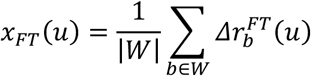

and

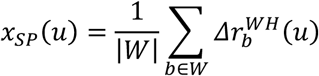

where *u* indexes units, *W* denotes the set of bins in the sliding window, and *b* indexes individual bins. A paired t-test was performed across units on the within-window difference *x_FT(u) − x_WH(u)*. For each task and alignment window, p-values from all sliding windows were corrected using the Benjamini–Hochberg FDR procedure (α = 0.05), yielding a Boolean significance mask over time.

#### Model dominance per unit

For each unit *u* and analysis window *w*, the modulation index (MI) was computed to quantify the relative dominance of semantic versus phonological encoding. The MI is defined as the difference in permutation-corrected correlations between the fastText (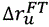) and Whisper models (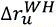):

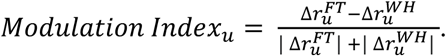

Positive values of the *Modulation Index*_*u*_ indicate semantic dominance, whereas negative values indicate phonological dominance.

#### Temporal window of interest

The modulation index was averaged within a prespecified temporal window (100–1000 ms after target onset):

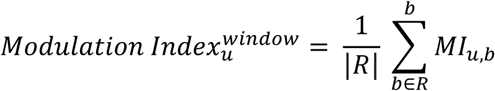

where *R* denotes the set of bins within the temporal window, and *b* indexes individual bins. These unit-wise window-averaged MI values were then averaged across units within each region and compared across tasks using a linear mixed-effects model with per-unit random intercepts:

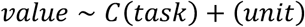

From the fitted model, pairwise contrasts between tasks on the modulation index were computed, and two-sided p-values were obtained. Within each window panel, p-values across the planned contrasts were adjusted using the Benjamini–Hochberg FDR procedure (p < 0.05); significant pairs are marked with asterisks. Outliers were clipped, and axis limits were scaled to percentiles to maintain readability.

#### Unit-wise significance

For each datapoint of unit × model × window, one-sided (“greater”) tests were performed within the prespecified window of interest in the target epoch (100 – 1000 ms after target onset). Within the ROI, mean correlations were computed in sliding sub-windows of width 150 ms and step 150 ms. For each sub-window *j*, the observed statistic was:

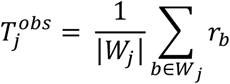

The same statistic was computed for each permutation k = 1, …, B, yielding 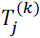. An empirical one-sided p-value was computed per sub-window as:

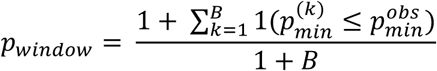

To correct for testing across the six sub-windows, a permutation-based min-p procedure (Westfall & Young) was applied: the observed minimum p-value across sub-windows was compared against the null distribution of the minimum p-value, obtained by computing, for each permutation *k*, the minimum p-value across all sub-windows. This yielded one familywise-error-rate-corrected p-value per unit × model, reflecting the best-responding sub-window while accounting for the implicit multiple comparisons. These corrected p-values were then submitted to BH-FDR correction (α = 0.05) within each session × model. Units were deemed significant when their FDR-adjusted *q*-value satisfied *q* ≤ 0.05. Within each session, units were categorised based on model-specific significance: semantics-only (significant for fastText but not Whisper), phonology-only (significant for Whisper but not fastText), significant for both, or neither. The “both” category reflects units for which the intersection-union test statistic, defined as max(*p*_*FT*_, *p*_*WH*_), itself survived BH-FDR correction within the same session × model scope.

### Population structure and representational similarity

The same trial sampling procedure as in the decoding and regression analysis were applied; i.e., trials were balanced per item (minimum six trials per item, irrespective of accuracy). Spike trains were discretized into 10-ms bins and convolved with a Gaussian kernel (150-ms standard deviation), yielding smooth peri-stimulus time histograms (PSTHs) as continuous firing rate estimates.

#### Demixed principal component analysis

We used demixed principal component analysis (dPCA)^44^, a dimensionality reduction method that extends principal component analysis (PCA) by decomposing neuronal variance into marginalizations aligned with task variables. Population activity was projected onto condition-averaged PSTHs and contributions from different task parameters were demixed. In this setting, neuronal variance was decomposed into three types of marginalizations: (i) time-dependent (t) marginalizations, capturing variance shared across items but varying systematically with trial time (e.g. attention, motor preparation, or task engagement); (ii) stimulus-dependent (s) marginalizations, capturing variance explained by differences between items; and (iii) stimulus–time interaction (st) marginalizations, capturing item-specific dynamics evolving over time. Fifteen components were extracted for each marginalization and method-intrinsic, automatic regularization was employed to prevent overfitting.

Following the authors’ guidelines, a minimum of six valid trials per session and at least 100 simultaneously recorded neurons per region and item subset were required to ensure robust demixing. As a result, ANG was excluded from these analyses. Data from multiple sessions were pooled whenever the same item groups were shared (i. e., subset of items), ensuring that all combinations of task parameters were represented and shared across instances of dPCA. These measures ensured that the demixed variance reflected stable and reproducible coding structure rather than session-specific variability. To quantify the extent to which the neuronal activity was captured by the demixed components, we computed the explained variance (EV). We followed the reconstruction-based definition implemented in the reference MATLAB code^63^. For each set of components, reconstructed PSTHs were obtained as

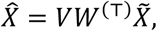

where 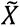 is the per-neuron centered data matrix (PSTHs with mean activity across conditions removed), *V* the encoder, and *W* the decoder. The explained variance was then defined as

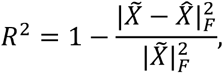

with | ⋅ |_*F*_ denoting the Frobenius norm. To partition variance into marginalizations (*ɸ*), the same procedure was applied to ANOVA-style decompositions 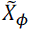, ensuring that contributions of time, item, and their interaction summed to the total variance.

We further computed time-resolved explained variance by repeating this calculation for each time point using per-time denominators 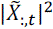. This yielded temporal trajectories of explained variance for each marginalization, revealing how different components contributed to population variance over the course of the trial. The analysis tracked changes in item coding, temporal dynamics, and their interactions at sub-second timescale, providing insights into the temporal structure of language processing in cortical populations. We then averaged explained variance values across all sessions for each brain region and marginalization type.

#### Implementation choices

We implemented the reconstruction-based EV calculation in Python, adapting the official MATLAB implementation. This was necessary because the available Python dPCA package reports projection energy along decoder axes rather than reconstruction-based variance. While equivalent for PCA, this metric can overestimate variance in dPCA, where components are not orthogonal, and can even yield cumulative values exceeding 100 %. For consistency with the original definition and published figures, we therefore used the reconstruction-based method, reporting fractions of PSTH variance (rather than PSTH plus noise variance).

#### Stimulus-related reconstructions

To focus on item-specific coding, we reconstructed neuronal activity from only the stimulus and stimulus-time interaction components:

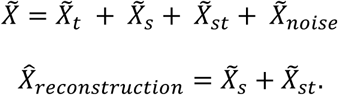

### Population vector analysis

To examine how neuronal populations represent different items, we analysed the geometric organisation of reconstructed neuronal stimulus representations in population activity (state) space. Splitting the trial into 100-ms bins, we computed population vectors (PVs) as the mean firing rate of each neuron in each bin. Differences between PVs corresponding to matched time points across items quantified representational dissimilarity. We quantified these differences using Euclidean distance in state space, as it captures both magnitude and directional differences between population states, providing a more geometrically faithful measure of separation than correlations. To obtain a robust measure of regional dynamics, distances were further averaged across item subsets, and word combinations. This provided an intuitive summary of general trends for each region and task, while preserving the metric’s sensitivity to item-specific divergence within trials. Distances were normalised by averaging within each region and task, which may overemphasise the contribution of individual conditions (for example, repetition in SMG).

#### Psycholinguistic representational similarity analysis

To relate neuronal distances to linguistic structure, psycholinguistic distances between words were quantified. Word similarity was defined using multiple measures: (i) phonological and semantic model embeddings (see phonological and semantic feature encoding) and (ii) Levenshtein edit distance for phonological similarity. High-dimensional embeddings (Whisper, fastText) were reduced with principal component analysis (PCA; n_components = 20) before computing cosine similarity across word combinations. While embeddings capture rich structure, clustering structure in phonological embeddings degraded when calculating the difference vector across items in embedding space. For semantic embeddings, clustering remained plausible. To address this, phonological edit distance (ii) was included as a complementary low-dimensional measure.

To assess the relationship between neuronal and linguistic distances for each task (*e*) and region (*r*), paired generalised linear models (GLMs) were fitted at every time bin (*t*). Separate models were run for each measure to avoid confounds between potentially correlated predictors. The semantic model used the cosine similarity of the 20-dimensional fastText embedding projections; the paired Levenshtein model used the scalar phonological distance. Fold-wise train–test splits (80/20 split, n = 100 iterations) produced out-of-sample coefficients of determination (R²). Importantly, GLMs allowed us to test whether linguistic structure could predict neuronal distances, rather than merely correlate with them. Randomly label-shuffled embeddings served as a control. Because demixed principal component analysis (dPCA) projects onto mean trial activity and phonological embeddings also contain speaker information, these embeddings were averaged across speakers. The difference in R² values between each predictor and its shuffled control was reported.

#### Bin-wise significance

To assess which time bins favored fastText over Levenshtein, we formed paired differences per iteration,

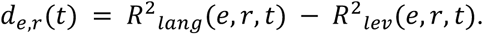

Two-sided paired Student’s t-tests compared *d*_*e*,*r*_(*t*) against zero. Resulting p-values were Benjamini-Hochberg corrected within each phase, and bins with *q* < 0.05 were flagged. We focused on the target epoch, reduced each fold to the window t ∈ [0.1, 1.0] after alignment, and computed the difference between fastText and Levenshtein per fold as:

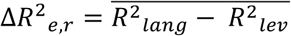

where the bar denotes the temporal average across the ROI. Taskwise means *μ*_*e*,*r*_) and standard errors were shown. Pairwise contrasts between experiments were quantified with Welch t-tests on the iteration distributions (*ΔR*^2^_*e*,*r*_). Resulting *p*-values were FDR-adjusted across the three pairs and rendered as bracket annotations (if *q* < 0.05).

## Extended Data

**Extended Data Fig. 1.**
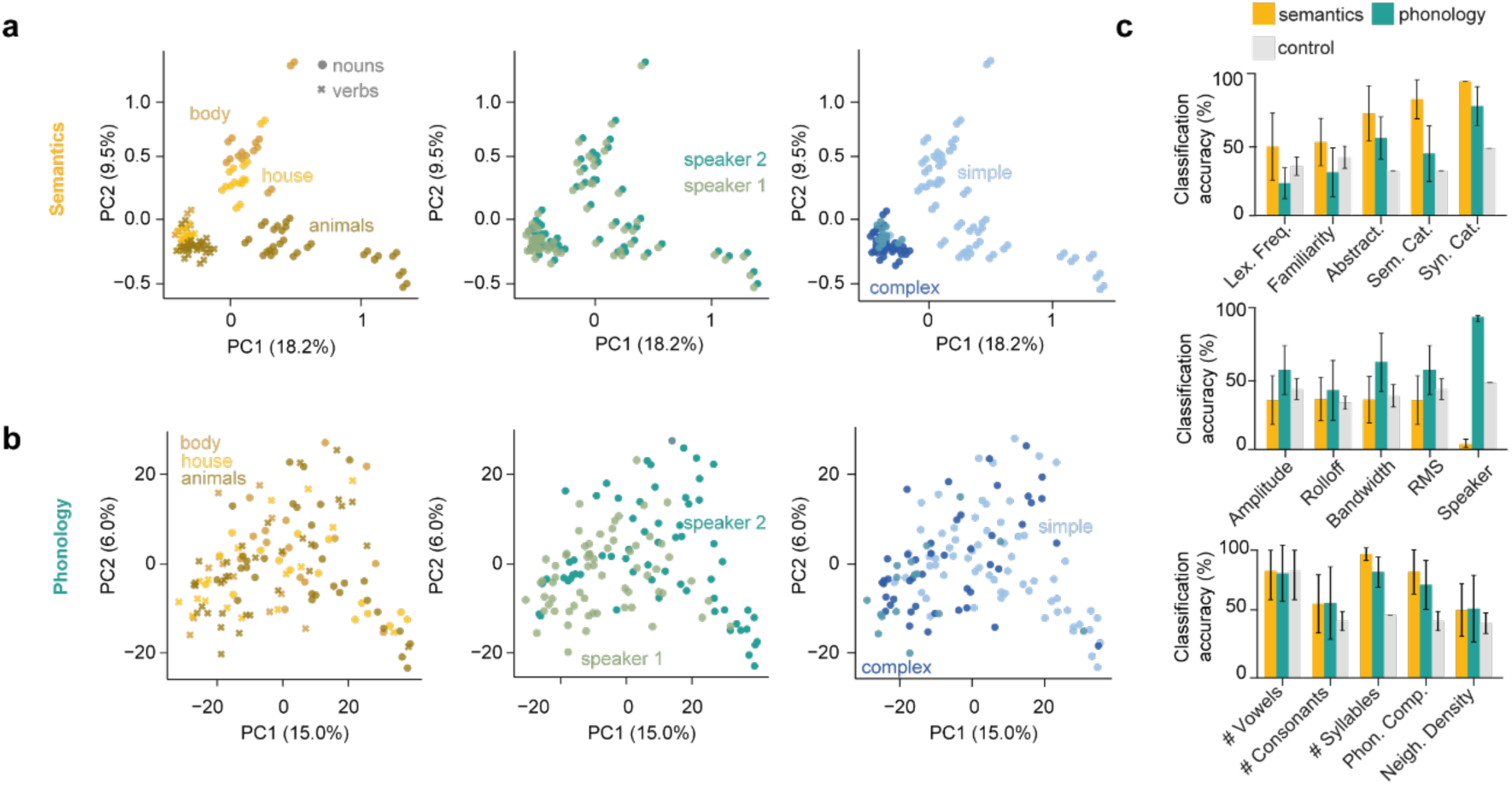
Semantic and phonological model embeddings. **a-b**, Embedding spaces for (**a**) the semantic model (fastText) and (**b**) the phonological model (Whisper). Each word was given to the models either as a text token or as an audio recording. Because two speakers produced the audio stimuli, each word appears twice in the phonological embedding space, while its text representation is unique. Embeddings are projected into two dimensions using PCA (with explained variance by PC shown in brackets). The semantic embeddings cluster by semantic category (left column), while the phonological embeddings primarily separate by acoustic properties, most clearly speaker identity (middle column). Both models also reflect phonological structure (right column). **c**, Balanced accuracy of a logistic regression classifier trained to predict lexical (lexical frequency, familiarity, abstractness, semantic category, syntactic category), acoustic (mean amplitude (discretized), mean rolloff (discretized), mean spectral bandwidth (discretized), root mean square (discretized), speaker identity), and sublexical features (number of vowels, number of consonants, number of syllables, phonological complexity, neighborhood density) from either semantic or phonological embeddings. Model accuracy was evaluated using 10-fold cross-validation and compared against a dummy baseline (most frequent class strategy).

**Extended Data Fig. 2.**
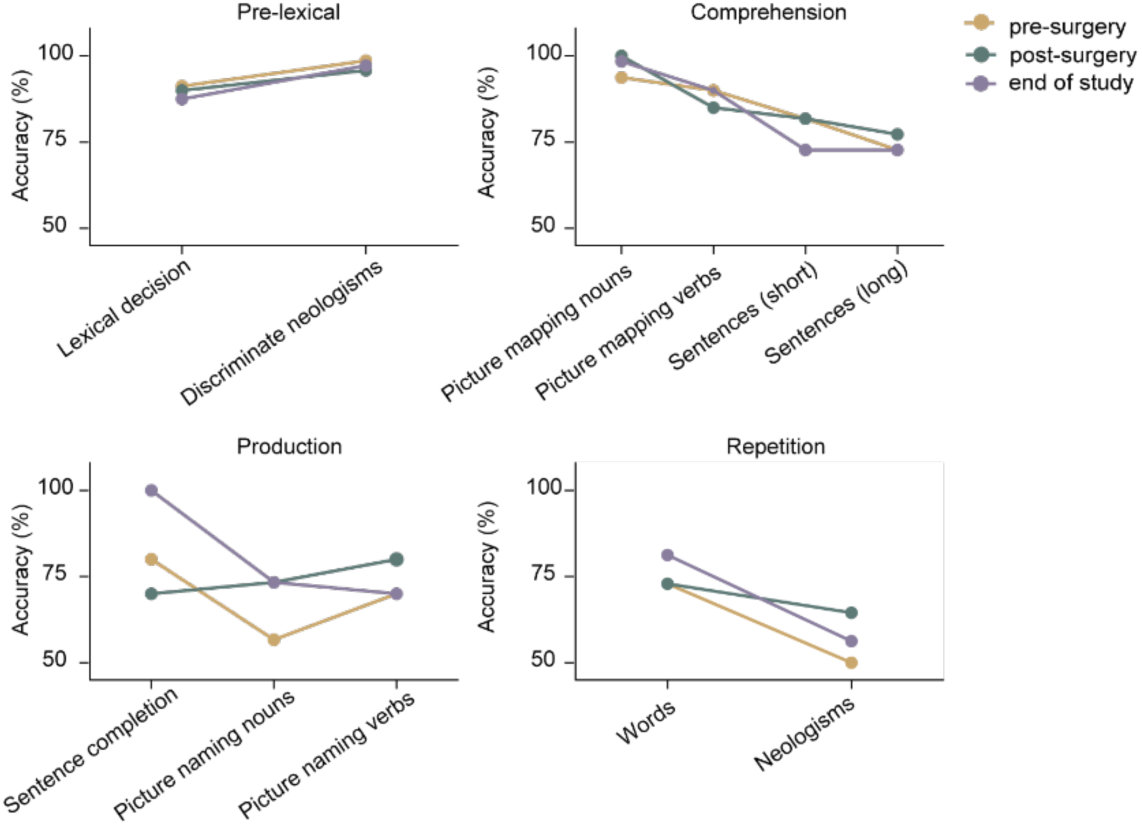
Language impairment profile. Language impairment profile assessed using standardized German language tests at three timepoints: one day prior to implantation (pre-surgery), two weeks post-surgery, and at end of study.

**Extended Data Figure 3.**
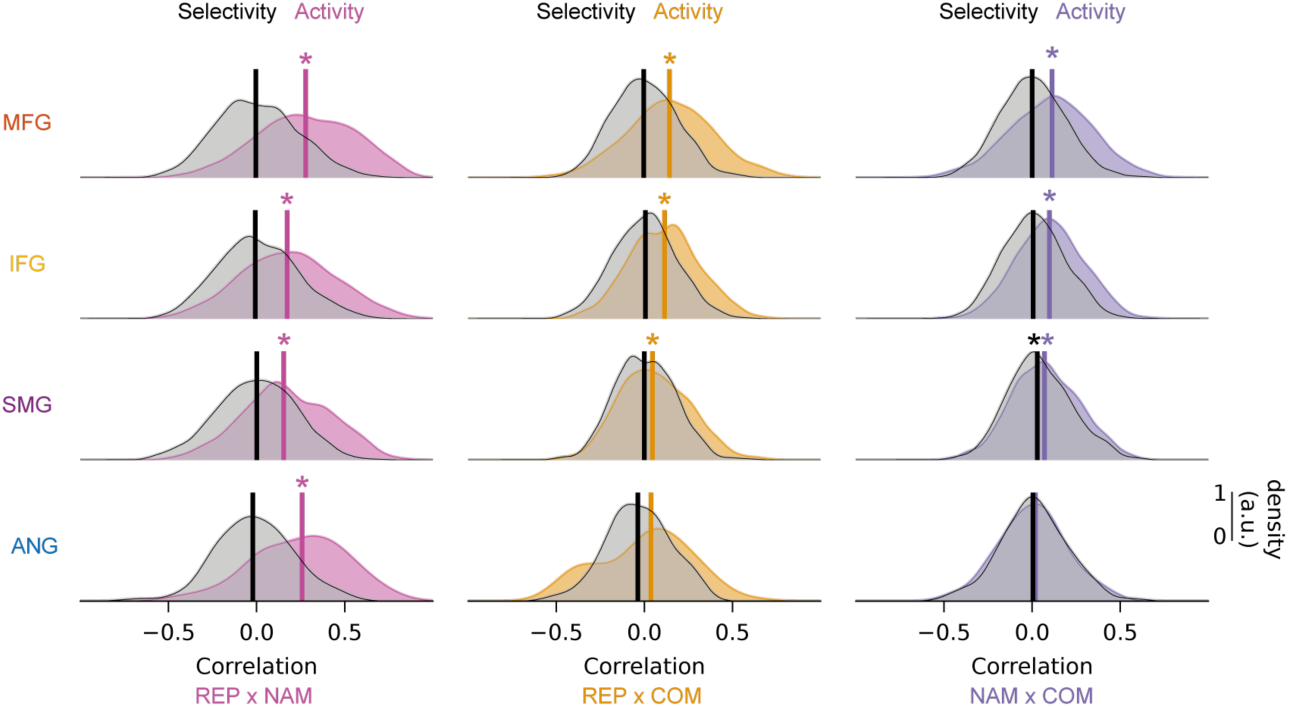
Activity and item selectivity correlation analysis. Density distributions of cross-task, unit-wise Pearson’s correlations of spiking activity (coloured) and item selectivity (black) for each region. Asterisks indicate distributions significantly greater than zero (one sample t-test, p < 0.001).

**Extended Data Figure 4.**
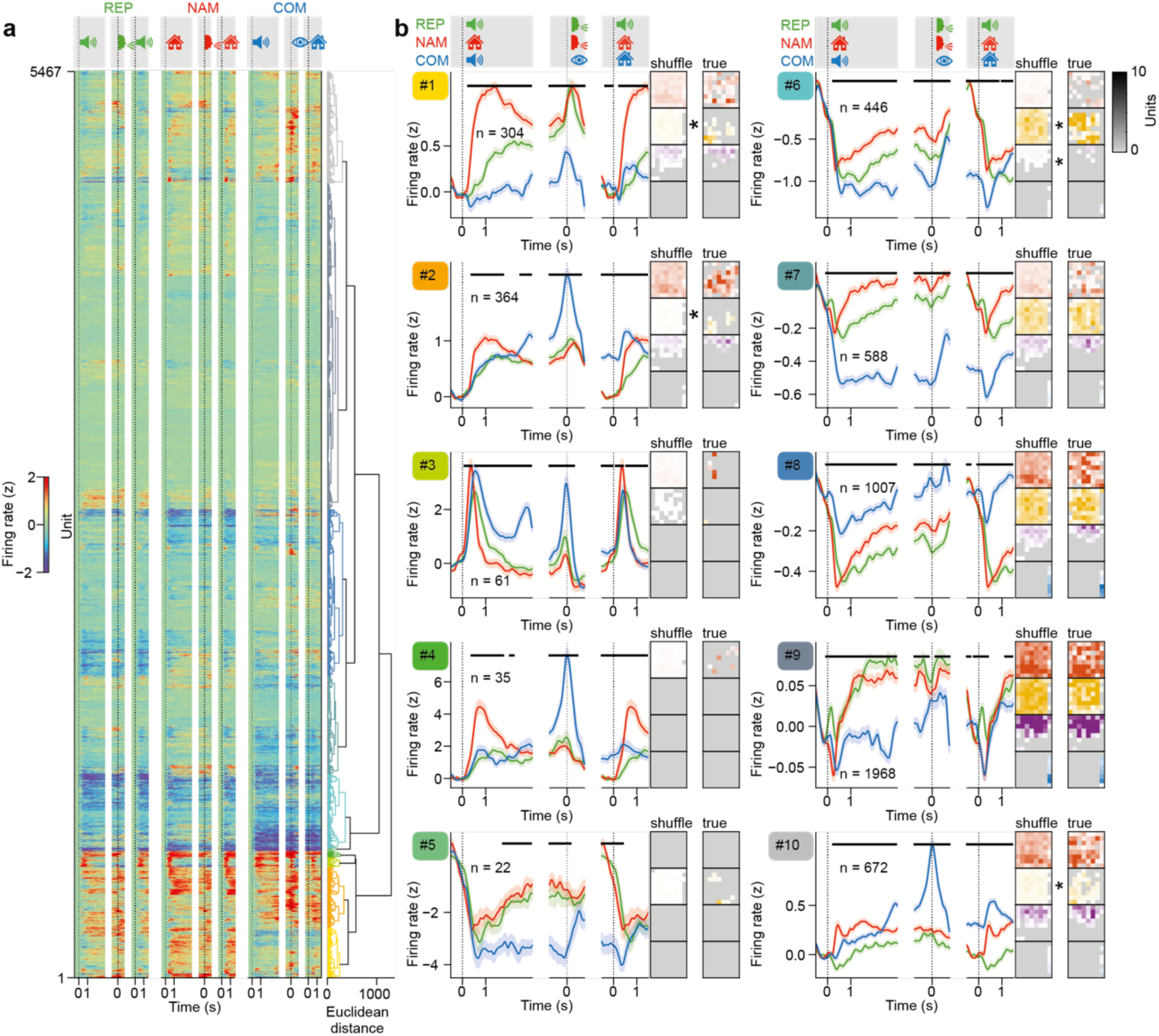
Clustering of activity profiles. **a**, Hierarchical clustering of spiking activity, as the z-scored trial-averaged responses to each task, performed across areas. Trials were randomly subsampled and balanced to minimally seven correct trials per item for minimally three items, and units with zero variance in baseline firing rate were excluded. The dendrogram colors indicate the cluster identity. **b**, Overview of all activity clusters, showing cluster-wise unit-averaged activity, split by task. Horizontal bars indicate periods with significant differences between tasks (ANOVA, p < 0.01). Error bands, SEM across units. Analyses were restricted to sessions in which all three language tasks were completed. Array columns show the spatial distribution of units across the four arrays. The left column shows the distribution based on session-wise shuffled cluster labels, while the right shows the true distribution. Asterisks indicate clusters that are significantly more spatially organised than the null distribution (permutation test, 99^th^ percentile).

**Extended Data Figure 5.**
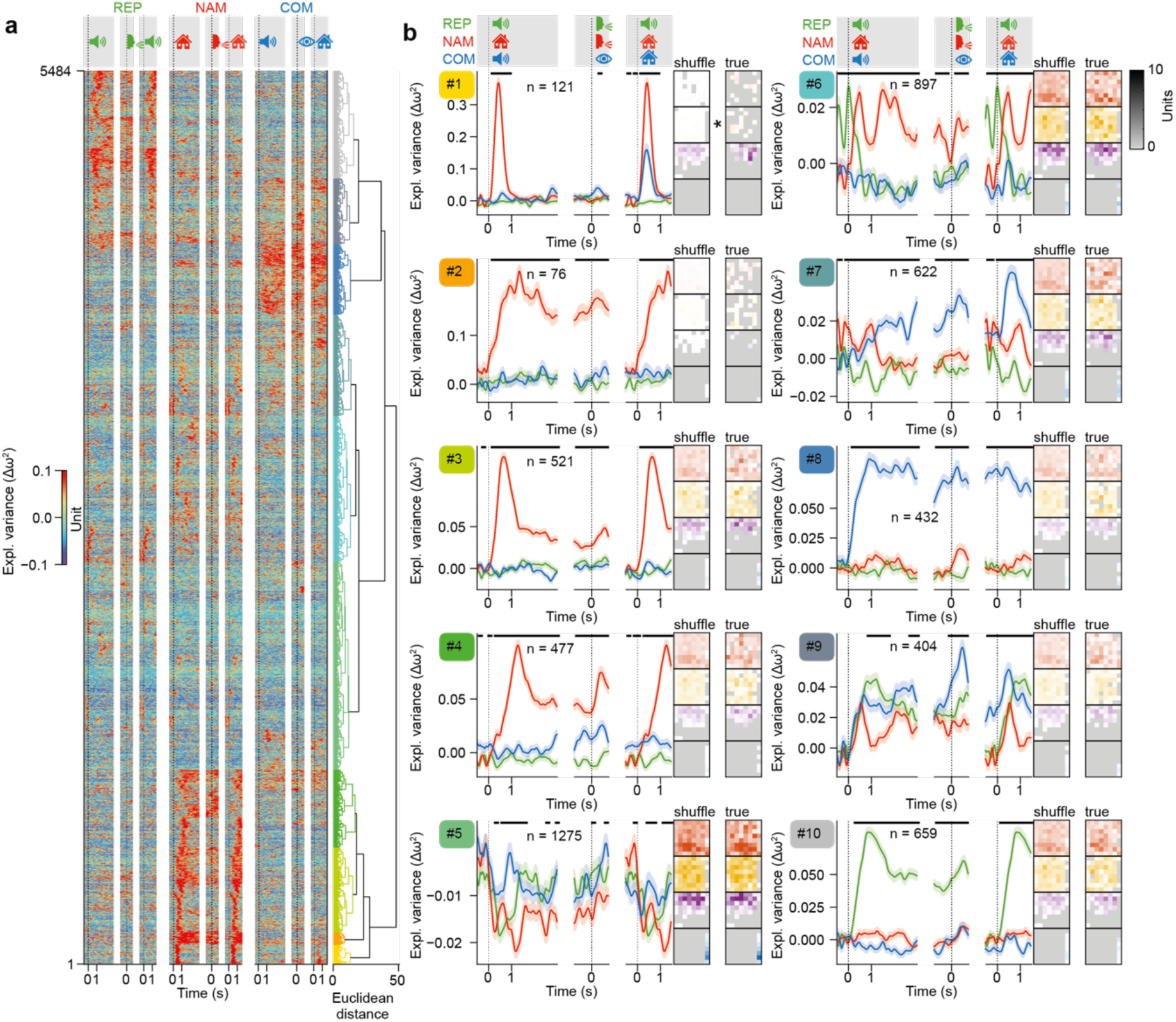
Clustering of item selectivity profiles. **a**, Hierarchical clustering of item selectivity, performed across areas. Trials were randomly subsampled and balanced to minimally seven correct trials per item for minimally three items, and units with zero variance in baseline firing rate were excluded. The dendrogram colors indicate the cluster identity. **b**, Overview of all item selectivity clusters, showing cluster-wise unit-averaged item selectivity, split by task with shuffled control subtracted (Δω²). Horizontal bars indicate periods with significant differences between tasks (ANOVA, p < 0.01). Error bands, SEM across units. Analyses were restricted to sessions in which all three language tasks were completed. Array columns show the spatial distribution of units across the four arrays. The left column shows the distribution based on session-wise shuffled cluster labels, while the right shows the true distribution. Asterisks indicate clusters that are significantly more spatially organised than the null distribution (permutation test, 99^th^ percentile).

**Extended Data Figure 6.**
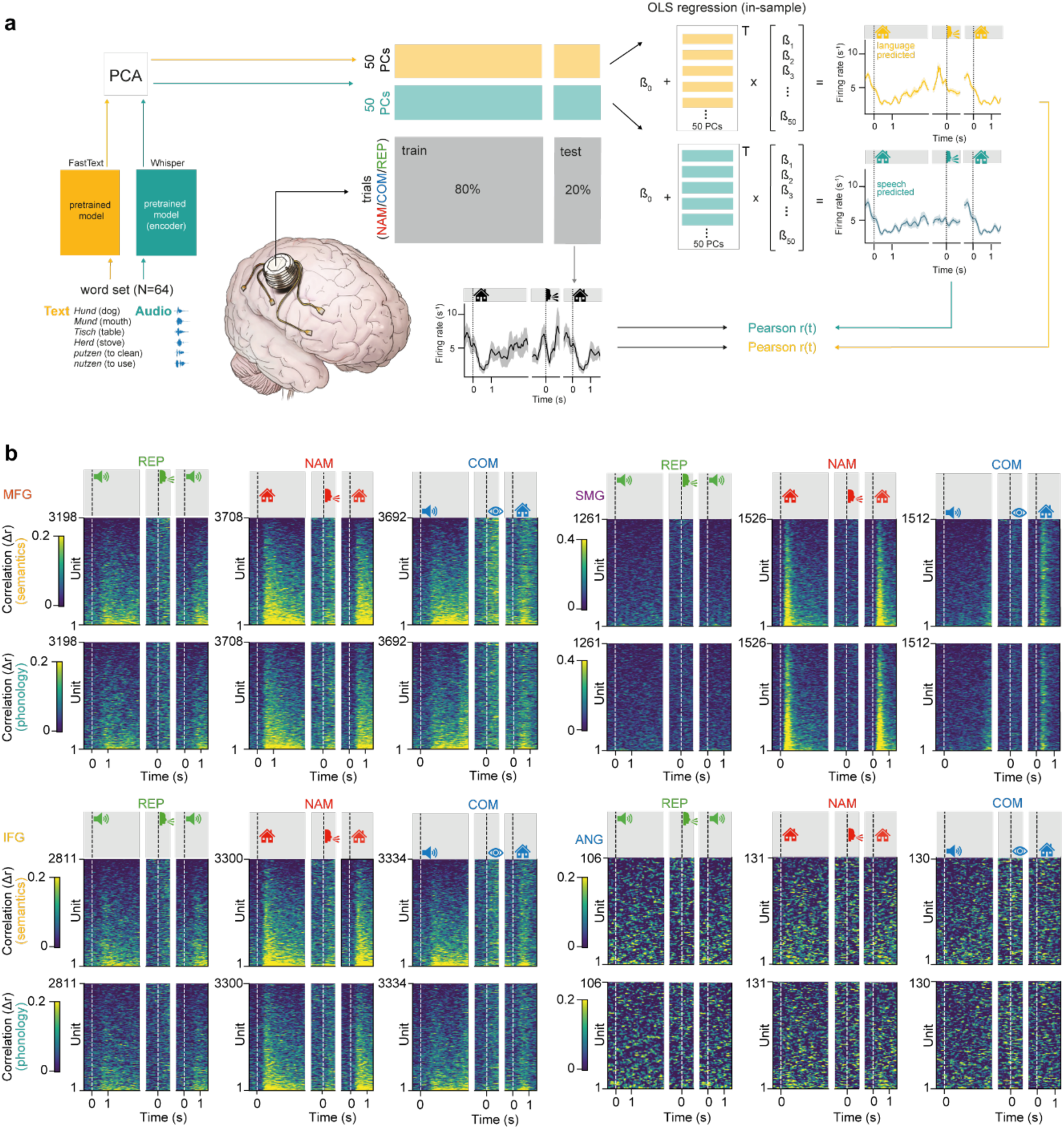
Unit-wise linear regression using model embeddings. **a**, Schematic of the time-resolved, unit-wise linear regression logic using semantic and phonological embedding components to predict firing rates of individual units at each electrode array. For each unit and window, models were trained on 80% of the trials firing rates were predicted on the held-out 20%. Model encoding was evaluated by correlating true firing rates with each model prediction at each time bin. **b**, Delta correlations (Pearson correlation between predicted and observed firing rates, computed on permutation-subtracted unit traces) for each unit for the semantic (top) and phonological (bottom) model across tasks. Units yielding NaN correlations due to zero variance in either trace were excluded; remaining units are sorted by their mean semantic correlation in the sample epoch. Unit counts are larger than in **Figs. 3, 4** and **6** as no co-recording requirement was imposed.

**Extended Data Figure 7.**
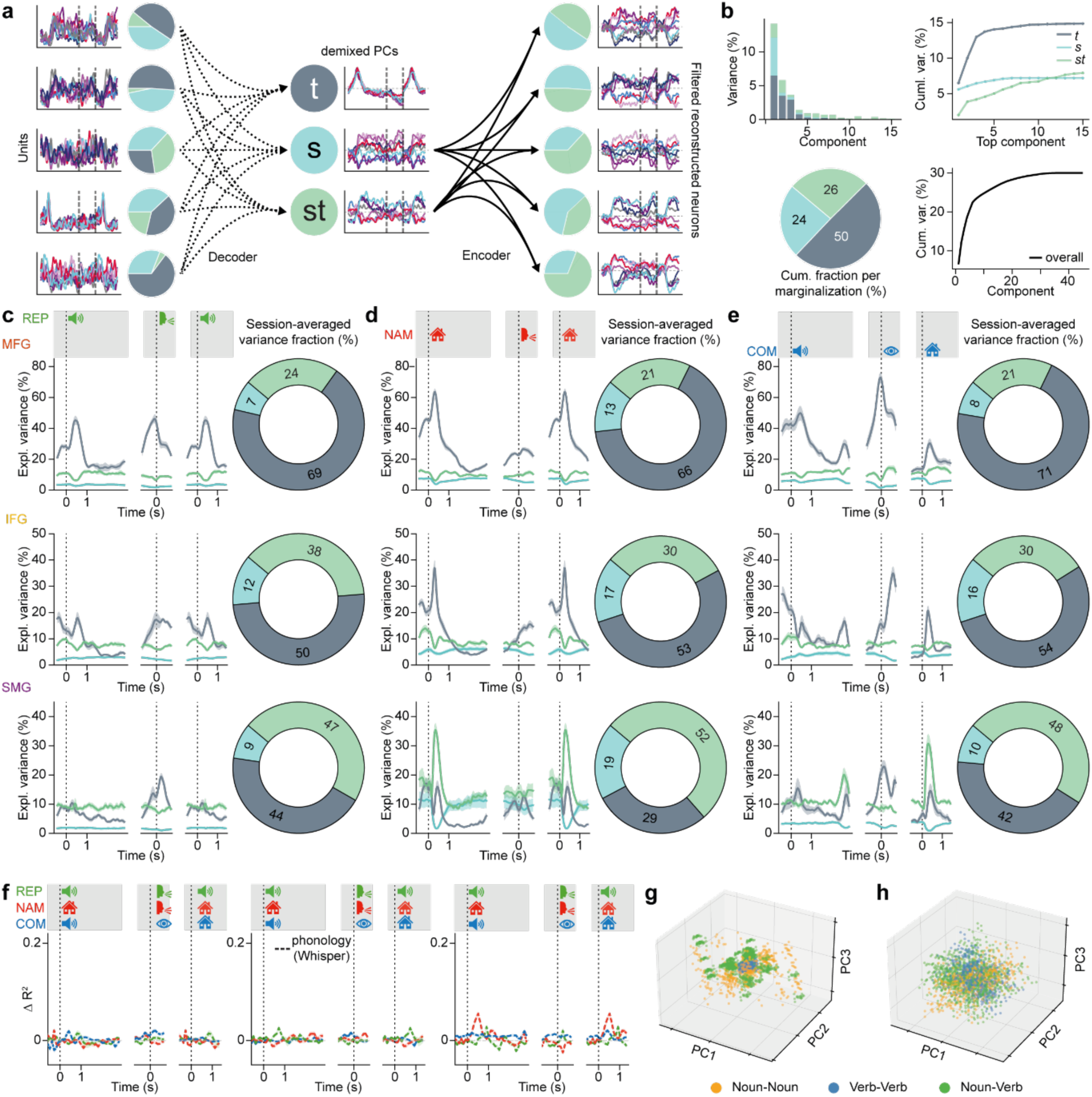
dPCA decomposition, encoding analyses, and semantic structure. **a**, Schematic of the dPCA “decoder–encoder” workflow. Trial-averaged spike trains were demixed into time (t), stimulus (s), and stimulus–time interaction (st) components. Filtered reconstructions, using only the s and st marginalizations, are projected back into neuron space to recover condition-specific firing trajectories. **b,** Example item subset (MFG during naming). Top: variance explained by individual components (left) and cumulative curves for each marginalization (right). Bottom: fraction of total variance captured by t, s, and st (left) and cumulative variance as a function of the number of total components (right). **c**–**e** Time-resolved explained variance in MFG, IFG, and SMG (top to bottom) across tasks (REP, NAM, COM; left to right). Lines indicate the across-combination mean, shaded bands show SEM of individual word subsets. Accompanying pie charts report the time-averaged variance fraction for each marginalization (t, s, st). **f**, Time-resolved encoding strength of GLMs for phonological (Whisper) embeddings in a representational similarity analysis. Panels show the difference in test-set coefficient of determination (R²) for generalized linear models predicting firing rate differences in MFG, IFG, and SMG (left to right). Models use pairwise word differences of speech-embedding features (cosine similarity in a 20-dimensional PCA space). The plotted quantity is feature-based R² minus shuffled-control R². Shaded bands denote SEM across cross-validated folds (n = 100; 80/20 split). Vertical bands mark sample, response, and target alignment windows. **g,** Semantic structure of pair-wise differences in embedding spaces. Language embeddings preserve category structure after dimensionality reduction; shown are PC1–PC3 projections of pairwise embedding distances stratified by category matches (within-category distances are smaller, indicating compact clusters). **h,** Similar to (**g**), differences in phonological (Whisper) embeddings analysed identically show weaker clustering, indicating reduced semantic separability.

**Supplementary Table S1.**
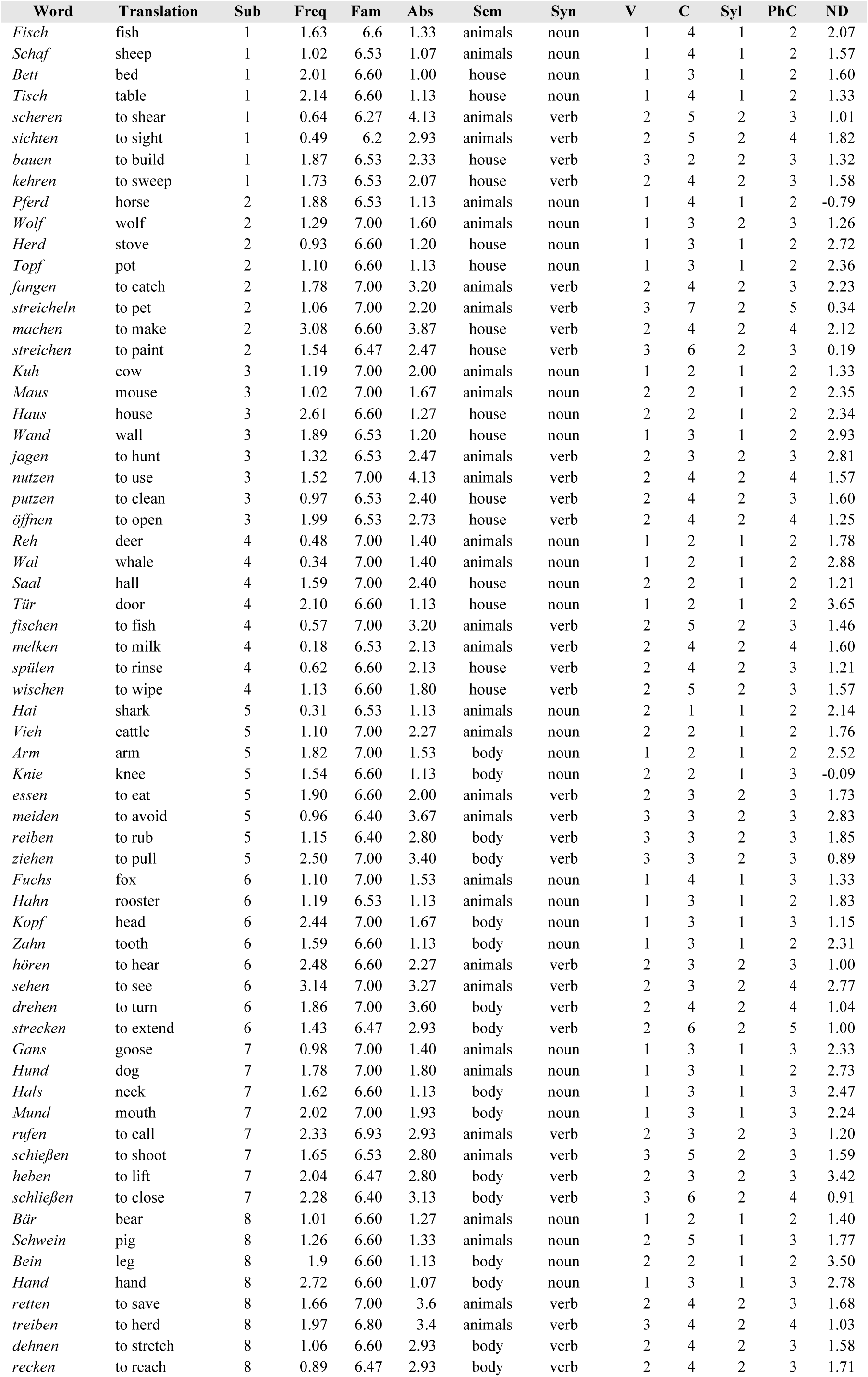
Lexical and phonological properties of the word set. The table lists lexical, semantic and phonological characteristics of the 64 German words used across all experimental tasks. Items were organised into eight subsets and belonged to three semantic categories (animals, household items and body parts) and two syntactic categories (nouns and verbs). Lexical frequency values (Freq) correspond to log word frequencies derived from the dlexDB lexical database for German. Phonological neighbourhood density (ND) values were also obtained from dlexDB. Familiarity (Fam) and abstractness (Abs) were obtained in a separate rating study in which 30 unimpaired native speakers of German rated the stimuli using Likert-scale questionnaires. Phonological descriptors include the number of vowels (V), number of consonants (C), number of syllables (Syl) and a phonological complexity score (PhC). Abbreviations: Trans, translation, Sub, experimental subset, Freq, lexical frequency (log frequency, dlexDB); Fam, familiarity rating; Abs, abstractness rating; SemCat, semantic category; SynCat, syntactic category; V, number of vowels; C, number of consonants; Syl, number of syllables; PhC, phonological complexity; ND, phonological neighbourhood density (dlexDB).

## Supplementary Material

Supplementary Information is available for this paper.

## Data Availability

The data that support the findings from this work are available upon reasonable request from the corresponding author. The data are not publicly available due to them containing information that could compromise research participant privacy.

## Code Availability

The custom MATLAB and Python scripts used for data analysis are available upon reasonable request from the corresponding author.

## Acknowledgements

We are grateful to our participant M.B. and her family for their pioneering spirit and extraordinary commitment, without which this work would not have been possible. We thank Franziska Schönweitz, Victoria Hohendorf and Christoph Penzkofer for creating visual and auditory stimuli, Johannes Hedemann for assistance with 3D brain reconstruction, and Leonie Kram, Sandro Krieg and Afra Wohlschläger for support with patient screening.

This work was supported by the TUM Innovation Network for Neurotechnology in Mental Health (Neurotech) to J.Ge., J.Gj., B.M and S.N.J., ERC STG 758032 (Memcircuit) to S.N.J. and ERC COG 101170179 (Rhetorical) to S.N.J.

## Contributions

Conceptualization: L.S., L.M.H., B.M., S.N.J.

Methodology: L.S., L.M.H., A.W., J.Ge., B.M., S.N.J.

Software: L.M.H., L.S., F.W., M.E., H.C., G.A., P.F.

Formal Analysis: L.M.H., L.S., F.W., M.E.

Investigation: L.S., L.M.H.

Resources: A.U., V.M.E., A.W., J.Ge., B.M., S.N.J.

Data Curation: L.M.H., L.S., H.C., G.A., P.F.

Writing - Original Draft: L.S.

Writing - Review & Editing: L.S., L.M.H., S.N.J.

Visualization: L.M.H., L.S., F.W., M.E.

Supervision: M.G.-W., J.Gj., B.M., S.N.J.

Project Administration: S.N.J.

Funding Acquisition: J.Gj., J.Ge., B.M., S.N.J.

## Competing interests

The authors declare no competing interests.

## Ethics declaration

This study was approved by the TUM University Hospital Institutional Review Board (IRB protocol number: 489/18 S-KK), and the participant provided written informed consent prior to participation.

## Notes

### Competing Interest Statement

The authors have declared no competing interest.

